# The white matter is a pro-differentiative microenvironment for glioblastoma

**DOI:** 10.1101/2020.11.14.379594

**Authors:** Lucy J. Brooks, Melanie P. Clements, Jemima J. Burden, Daniela Kocher, Luca Richards, Sara Castro Devesa, Megan Woodberry, Michael Ellis, Leila Zakka, Zane Jaunmuktane, Sebastian Brandner, Gillian Morrison, Steven M. Pollard, Peter B. Dirks, Samuel Marguerat, Simona Parrinello

## Abstract

Glioblastomas are hierarchically organised tumours driven by glioma stem cells that retain partial differentiation potential. Glioma stem cells are maintained in specialised microenvironments, but how they undergo lineage progression outside of these niches remains unclear. Here we identify the white matter as a differentiative niche for glioblastomas with oligodendrocyte lineage competency. Tumour cells in contact with white matter acquire pre-oligodendrocyte-like fate, resulting in decreased proliferation and invasion. Differentiation is a response to white matter injury, which is caused by tumour infiltration itself in a tumoursuppressive feedback loop. Mechanistically, tumour cell differentiation is driven by selective white matter upregulation of SOX10, a master regulator of normal oligodendrogenesis. SOX10 overexpression or treatment with myelination-promoting agents that upregulate endogenous SOX10, mimic this response, leading to white matter-independent pre-oligodendrocyte-like differentiation and tumour suppression *in vivo*. Thus, glioblastoma recapitulates an injury response and exploiting this latent programme may offer treatment opportunities for a subset of patients.

## Introduction

Glioblastoma (GBM) is the most common and malignant primary brain tumour (Stupp et al., 2005). Stark resistance to current treatments, which include maximal surgical resection, chemo- and radiotherapy, leads to tumour relapse in virtually all patients with a median survival of less than 15 months (Weathers and Gilbert, 2014). GBM initiation, growth and recurrence are thought to be rooted within a subpopulation of therapy-resistant tumour cells with properties of normal neural stem cells, termed glioma stem cells (GSCs) (Bao et al., 2006; Chen et al., 2012; Galli et al., 2004; Lathia et al., 2015; Singh et al., 2004).

Mounting evidence indicates that GSCs fuel tumourigenesis by recapitulating normal neural lineage hierarchies of self-renewal and generation of non-dividing progeny (Lan et al., 2017; Neftel et al., 2019; Park et al., 2017). Fate decisions of normal neural stem cells are tightly controlled by the microenvironment, which maintains stemness in specialised niches and directs differentiation along the appropriate lineages (Silva-Vargas et al., 2013). Similarly, it is well established that GSCs are maintained in perivascular and hypoxic regions of the tumour bulk (Calabrese et al., 2007; Hambardzumyan and Bergers, 2015; Li et al., 2009). In contrast, little is known about the mechanisms by which GSCs differentiate within tumours and whether pro-differentiative microenvironments exist. However, improved understanding of lineage progression is central to our understanding of the disease and may reveal novel strategies to suppress GBM growth and recurrence by directing GSC fate towards a more mature, non-dividing state. Consistent with this idea, manipulation of candidate developmental pathways to promote tumour cell differentiation into astrocyte and neuronal lineages has shown efficacy in preclinical models (Park et al., 2017; Piccirillo et al., 2006).

Diffuse infiltration into the normal brain parenchyma is a hallmark of GBM and underlies recurrence by precluding complete surgical resection (Cuddapah et al., 2014; Vehlow and Cordes, 2013). As GBM cells infiltrate away from the tumour bulk, they are confronted with new and heterogeneous microenvironments, which would be predicted to affect fate decisions (Molofsky et al., 2012). Indeed, it has been proposed that invading GSCs may lose stemness, but this remains controversial, with studies both in support and against this idea (Brooks and Parrinello, 2017; Hoelzinger et al., 2005; Molina et al., 2010; Piccirillo et al., 2009). A predominant route of infiltration is the white matter, which consists of bundles of myelinated axons, known as tracts, and represents approximately 60% of the brain (Blumenfeld, 2010; Cuddapah et al., 2014). GBMs frequently develop within white matter and spread throughout the brain using myelinated fibres as scaffolds for cell migration (Scherer, 1938). This includes spread to the contralateral hemisphere, which occurs exclusively along the white matter tracts of the corpus callosum (Cuddapah et al., 2014). Remarkably, the cellular and molecular mechanisms that underpin white matter invasion remain almost entirely unknown.

Here, we investigated the impact of the white matter microenvironment on tumour cell fate by profiling brain region-specific transcriptomes of GBM cells invaded into white and grey matter, alongside matched bulk cells. Unexpectedly, this analysis revealed that the white matter dominantly suppresses malignancy by directing GSC differentiation towards pre-oligodendrocyte-like fate. We show that this process recapitulates an injury response and can be harnessed therapeutically to suppress tumourigenesis in a subset of GBMs.

## Results

### Region-specific GBM transcriptomes

Despite the fundamental role of white matter in GBM biology, how this specialised microenvironment might modulate tumour cell behaviour remains unknown. To probe white matter phenotypes, we made use of a well characterised patient-derived GSC line with propensity to invade along white matter tracts (G144) (Pollard et al., 2009). GFP-labelled G144 cells were stereotactically injected into the striatum of immunocompromised mice and, upon development of clinically apparent disease, the tumour bulk (B) and margin regions extending into the corpus callosum (CC) as a region of white matter, and the striatum (ST) as a region of grey matter, were micro-dissected under fluorescence guidance (Figure 1A). Tumour tissue was dissociated to single cells and GFP^+^ tumour cells FACS-purified, pooled and processed for RNA-sequencing (RNA-seq). Bioinformatics analysis of human tumour reads revealed that margin cells are transcriptionally distinct from bulk cells, with invasive cells acquiring markers of lineage progression towards astrocytes, as well as downregulating proliferation signatures, which resulted in decreased EdU incorporation *in vivo* (Figure 1B, C, Figure S1A-E and Tables S1-4) (Cahoy et al., 2008; Zhang et al., 2014; Zhang et al., 2016). Strikingly, it also indicated that the transcriptomes of CC and ST tumour cells differed from one another (Figure 1B). Tumour cells invading into white matter selectively upregulated signatures of oligodendroglia, suggesting that myelinated regions may promote progression along the oligodendrocyte lineage (Figure 1C, Figure S1F and Tables S1-4). In agreement with this, CC cells upregulated a panel of oligodendrocyte lineage marker genes, including the transcription factor SOX10, a master regulator of oligodendrogenesis (Figure 1D) (Elbaz and Popko, 2019; Weider et al., 2013; Zhang et al., 2014). Notably, upregulation of myelin genes was mild and incomplete in the CC, indicative of partial differentiation, as expected from cancer cells (Azzarelli et al., 2018). To determine whether these gene expression changes also correlated with phenotypic changes, we characterised the differentiation response at the single cell level in two sets of complementary experiments. First, we dissociated G144 cells from the CC, ST and B of xenografts, seeded them acutely in mitogen-free media and assessed expression of SOX10, the pre-oligodendrocyte marker O4 and the myelinating oligodendrocyte marker myelin basic protein (MBP) by immunocytochemistry (Figure S1G, H). Preparations isolated from the CC had a strong increase in the proportion of cells positive for SOX10, the majority of which also expressed O4, but not MBP, confirming that transcriptomic signatures reflect changes in tumour cell fate. Second, we examined lineage marker expression within xenografts *in situ* by immunohistochemistry. GFP or the human-specific nuclear marker (NuMA) were used to label tumour cells and distinguish them from endogenous mouse glia (Figure S1I, J). The majority of tumour cells began to express high levels of SOX10 in white matter, including in myelinated fibres inside the tumour bulk (Figure 1E, F). SOX10^+^ tumour cells were significantly less proliferative than tumour cells that remained SOX10^−^ within the same region (Figure 1G, H) and occasionally expressed the immature oligodendrocyte markers CNP and CC1 (Figure S1 K,L), but were again negative for the mature marker MBP (not shown). To further assess the white matter specificity of this response, we implanted G144 cells directly into the upper layers of the cortical grey matter or in the corpus callosum and examined their differentiation using SOX10 induction as a read-out. Differentiated SOX10^+^/EdU^−^ tumour cells were only found in the CC, confirming that the white matter selectivey promotes partial GBM differentiation (Figure 1I, J). Thus, GBM cells differentiate to a pre-oligodendrocyte/immature pre-oligodendrocyte state in white matter, which is marked by induction of SOX10 expression.

**Figure 1.**
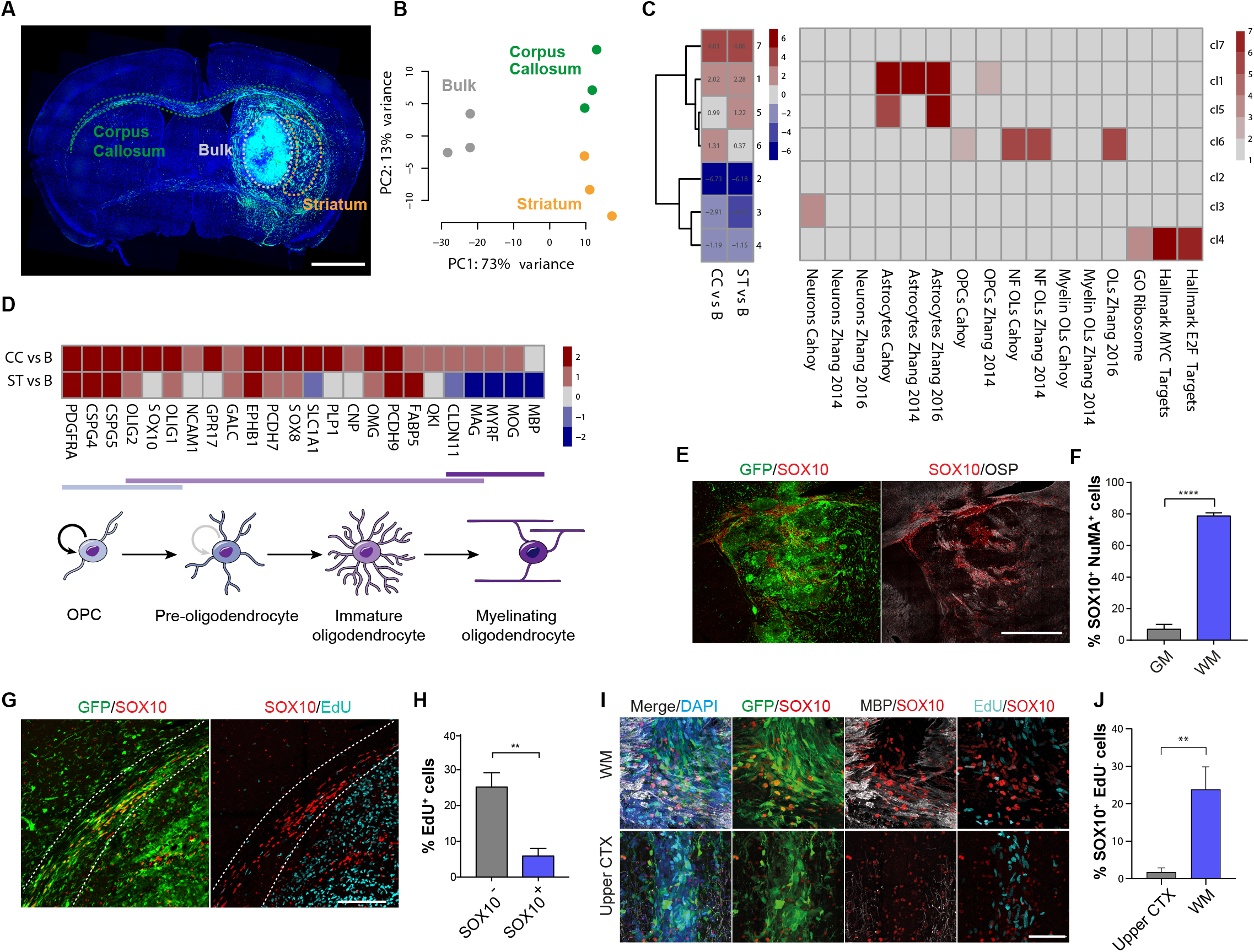
GBM cells invading into white matter acquire pre-oligodendrocyte fate. **A**, representative fluorescence image of a G144 xenograft collected for region-specific RNA-seq. Brain regions microdissected for FACS purification of GFP^+^ cells are shown, nuclei are counterstained with DAPI (blue). **B**, PCA plot of rlog-normalised DESeq2 expression scores for the resulting RNA-seq libraries. n=3 xenografts. **C**, left panel: K-means clustering of DESeq2 expression ratios (absolute log_2_ expression ratio > 0.58, p_adjust_ < 0.05) for tumour cells invading into the corpus callosum (CC) or striatum (ST) relative to the tumour bulk (B). Right panel: overlap of K-means cluster with gene signatures for OPC, Oligodendrocytic, Astrocytic and Neuronal lineages as well as proliferation signatures. Colours represent −log_10_ of enrichment adjusted p-values. **D**, top: DESeq2 log_2_ expression ratios of markers of the oligodendrocyte lineage organised by differentiation stage in tumour cells from CC and ST relative to B. bottom: schematic illustration of the maturation of oligodendrocyte lineage cells. **E,**representative immunofluorescence staining for SOX10 (red) and the myelin marker OSP (grey) of GFP-labelled G144 tumours (green). Scale bar=1mm. **F**, quantification of percentage of SOX10^+^ tumour cells in grey (GM) and white matter (WM). A minimum of 750 cells per xenograft were counted. Mean±SEM, n=3 xenografts ****p<0.0001. Unpaired two-tailed Student’s t test. **G**, representative EdU (turquoise) and SOX10 (red) staining and **H**, quantification of actively proliferating SOX10^+^ and SOX10^−^ GFP^+^ tumour cells in the corpus callosum. Scale bar=500μm. A minimum of 1000 cells per xenograft were counted. Mean±SEM, n=3 xenografts. *p<0.05 **I**, representative SOX10^+^ (red)/EdU^−^ (Turquoise) immunofluorescence staining and **J**, quantification of GFP^+^ tumour cells implanted in the white matter of the corpus callosum (top) or the grey matter of the upper cortex (bottom). A minimum of 140 cells per xenograft were counted. Scale bar=100μm Mean±SEM, n=3 xenografts **p<0.01. Unpaired two-tailed Student’s t test. See also Figure S1.

To determine the generality of these findings, we xenotransplanted five SOX10^+^ GSC lines (three of which were freshly dissociated from primary tumours) and examined their behaviour in white matter (Table S5). One line had lost white matter regulation of SOX10 induction, displaying homogeneous high SOX10 expression throughout the xenograft and was not analysed further (G564). In contrast, the other four lines were similar to G144 cells, with a somewhat variable, but consistent increase in the proportion of SOX10^+^ cells in white matter as compared to grey matter (Figure 2A, B and Figure S2A). In addition, as in G144 xenografts, SOX10^+^ cells proliferated significantly less than SOX10^−^ cells in white matter and partially progressed to CNP^+^ or CC1^+^ immature pre-oligodendrocyte cells (Figure 2C and Figures S1K, L and S2B). To determine whether the white matter-induced response also occurs in primary patient tumours, we took two complementary approaches. First, we selected three SOX10^+^ cases based on retained SOX10 expression in derivative GSC lines upon xenotransplantation (GCGR-L12, GL67 and GL23, Table S5). We found that in all cases the white matter contained SOX2^+^ tumour cells, which upregulated SOX10 and proliferated less than their SOX10^−^ counterparts, confirming that GSC behaviour in xenograft models reflects disease phenotypes (Figure 2D, E and Table S6) (Singh et al., 2004). SOX2 staining was specific to GBM cells, as the vast majority of endogenous SOX10^+^ OPCs were negative for SOX2 in areas of tumour infiltration (Figure S2C). Second, we selected 23 white matter-containing cases without prior knowledge of SOX10 status. Remarkably, we found that 15 cases contained at least a subset of SOX10^+^ cells in white matter, regardless of molecular characteristics (Figure 2F, S2D and Table S6). In addition, SOX10^+^ cells were lowly proliferative, progressed to a CNP^+^/CC1^+^/MBP^−^ immature pre-oligodendrocyte state and were restricted to MBP^+^ tumour regions (Figure 2F and Figure S2E-G). These experiments confirm that a subset of GBMs progress to pre-oligodendrocyte-like cells in white-matter.

**Figure 2.**
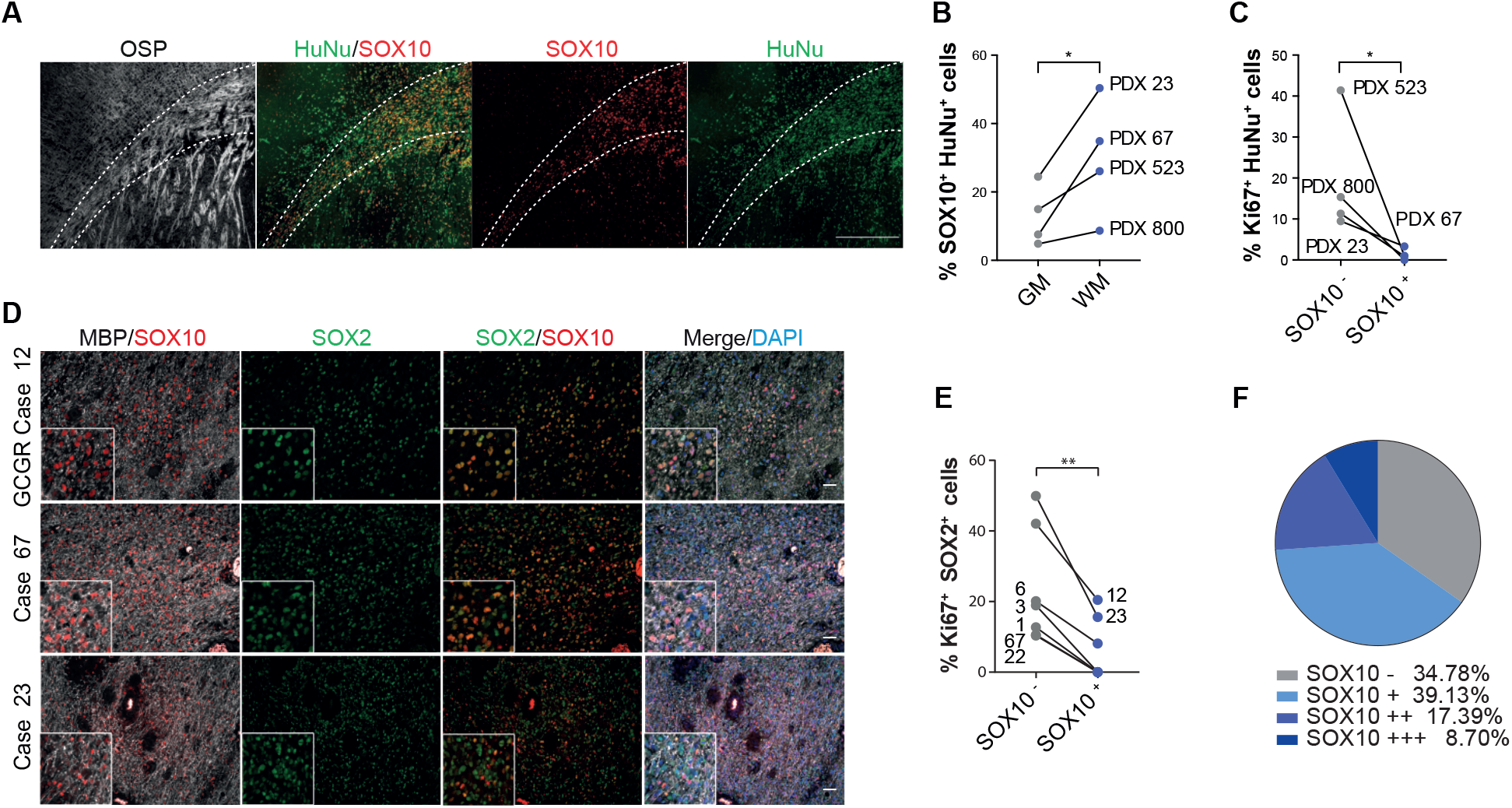
Differentiation in white matter is a general GBM response. **A**, representative immunofluorescence image of a SOX10^+^ patient-derived xenograft (PDX 23) stained for SOX10 (red), OSP (grey) and HuNu (green) to label all tumour cells. Scale bar=500μm. **B**, quantification of percentage of SOX10^+^/HuNu^+^ tumour cells in white and grey matter of indicated patient-derived xenografts. n=4 independent xenograft models. For each xenograft, a minimum of 550 cells was quantified across 2 independent ROIs selected within white or grey matter. *p<0.05, paired one-tailed Student’s t test. **C**, quantification of percentage of proliferating SOX10^+^ and SOX10^−^ tumour cells in the corpus callosum of indicated xenografts. For each xenograft, a minimum of 500 SOX10^+^ or SOX10^−^ cells was quantified across 2 independent ROIs, n=4 independent xenograft models. *p<0.05, paired one-tailed Student’s t-test. **D**, MBP (grey) and SOX10 (red) immunofluorescence staining of white matter regions of patient tumours. SOX2 (green) was used to identify tumour cells and distinguish them from resident glia. Cases shown are the original tumours from which lines GL23, GL67 and GCGR L12 used in this study have been isolated. Inset shows a close-up of the same image. Scale bar=50 μm. **E**, quantification of percentage of SOX2^+^/SOX10^+^ and SOX2^+^/SOX10^−^ tumour cells undergoing proliferation (Ki67^+^) in patient tumours. For each case, a minimum of 300 cells was quantified across 2 independent ROIs, n=7 cases. *p<0.05, paired one-tailed Student’s t-test. **F**, pie chart representation of SOX10 expression in 23 patient tumours. – denotes absence, and + to +++ increasing frequency of SOX10^+^ cells within the tumour. See also Figure S2.

### Disrupted white matter drives pre-oligodendrocyte differentiation

Next, we sought to understand what properties of white matter are differentiation-promoting. We noticed that SOX10 induction was most robust in areas where OSP staining appeared disrupted, suggesting that it might be linked to demyelination (Figure S1I). We therefore assessed myelin integrity in xenografts by fluoromyelin staining and by electron microscopy (EM). The proportion of SOX10^+^/Human nuclear antigen (HuNu)^+^ tumour cells was highest in heavily infiltrated, fluoromyelin negative, white matter areas, as compared to dye-positive, low tumour-density, myelin regions in four independent xenografts (Figure 3A, B and Figure S3A). A similar differentiation pattern was also found in patient tissue, where only few SOX2^+^ tumour cells upregulated SOX10 in intact myelin (Figure S3B). EM analysis of G144 xenografts further revealed severe axonal pathology and demyelination in highly infiltrated white matter regions, including axonal swelling and vacuolisation, presence of dark axons, myelin decompaction and a significant increase in g-ratios relative to contralateral normal brain (Figure 3C-I and Figure S3C). In contrast, g-ratios remained normal in the tumour-infiltrated striatal grey matter (Figure 3H), indicating that demyelination is specific to white matter. Consistent with this, activated microglia increased selectively in tumour-infiltrated white matter and endogenous oligodendrocytes were significantly decreased in the CC (Figure S3D-G). Furthermore, activated microglia with engulfed myelin debris was frequently observed in both xenografts (Video S1) and human samples, and colocalised with SOX10^+^ tumour cells (Figure S3H-J). Surprsingly, despite severe myelin dysfunction, the mice did not exhibit overt motor or cognitive deficits.

**Figure 3.**
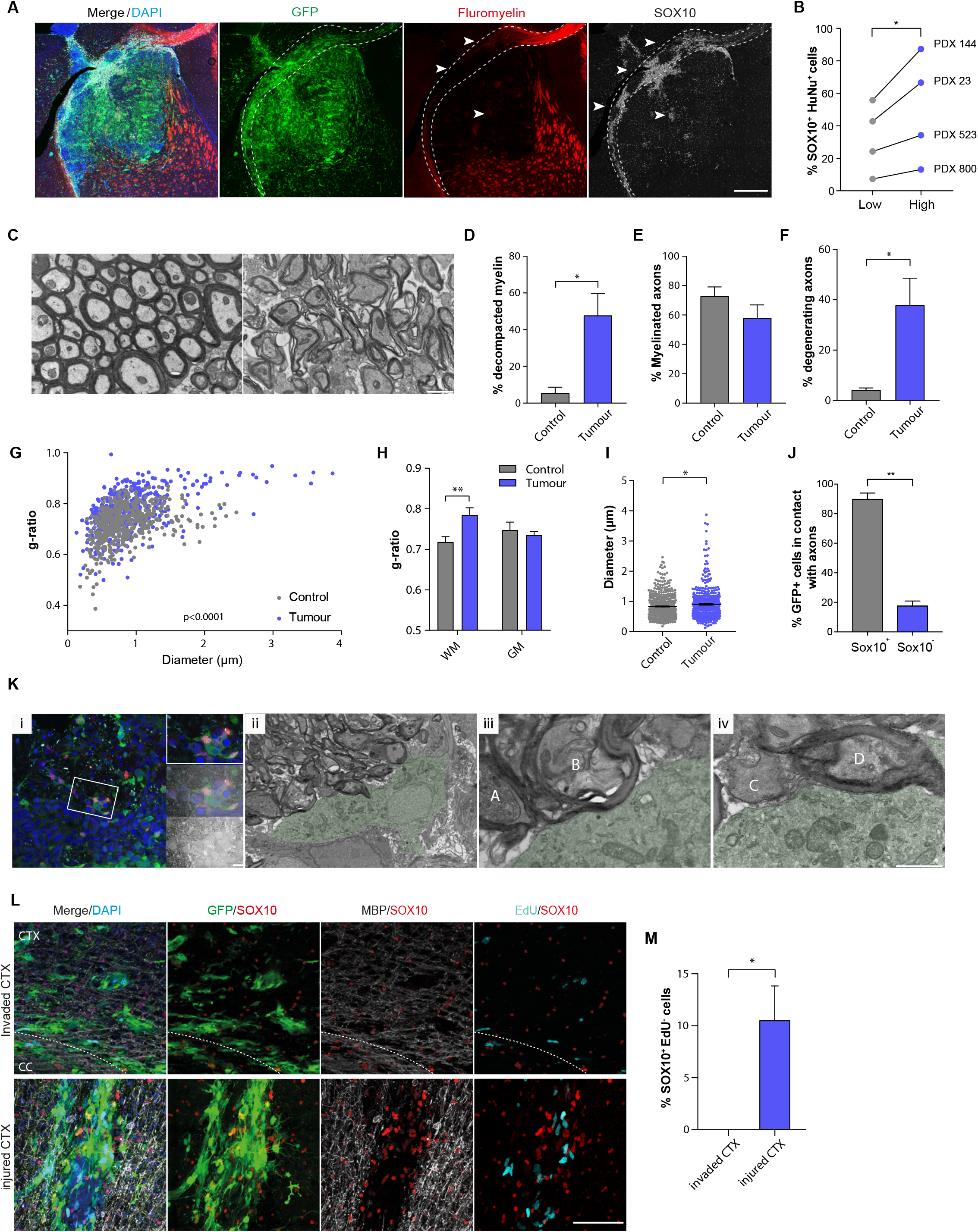
Exposure to disrupted myelin drives GBM progression down the oligodendrocyte lineage. **A**, SOX10 (grey), fluoromyelin (red) and DAPI (blue) immunofluorescence staining of GFP^+^ G144 xenografts. Arrowheads denote areas of demyelination. Scale bar=500μm. **B**, quantification of percentage of SOX10^+^ tumour cells in areas of high (blue) and low (grey) myelin disruption in the indicated xenograft models. A minimum of 120 cells was quantified across at least 6 ROIs selected within intact or disrupted white matter. n=4 xenografts. *p<0.05, paired one-tailed Student’s t test. **C**, Electron micrographs of myelin bundles in the ipsilateral (infiltrated) and contralateral (intact) striatum of G144 tumour brain. Scale bar=1μm. **D-I**, quantification of indicated axonal and myelin phenotypes in the EM data shown in b. n=3-4 xenografts. **p<0.01, *p<0.05, paired one-tailed Student’s t test, 2-way ANOVA or linear regression. **J**, quantification of percentage of GFP^+^ tumour cells in contact with axons within disrupted white matter. For each xenograft, a minimum of 300 SOX10^+^ or SOX10^−^ cells was quantified across 3 independent ROIs. Mean±SEM, n=3 tumours, **p<0.01. unpaired two-tailed Student’s t test. **K**, *i* Correlative light and electron micrograph (CLEM) depicting a GFP^+^(green), SOX10^+^ (red) pre-oligodendrocyte-like tumour cell directly interacting with multiple axons within white matter. EM micrographs are overlaid with corresponding confocal fluorescence images to conclusively identify tumour cells. *ii-iv* are magnifications of *i* highlighting interactions with decompacted myelin (A, B), a naked (C) and an intact myelinated axon (D Scale bar=10μm (i), 5μm (ii), 1μm (iii, iv). **L**, SOX10 (red), MBP (grey), EdU (turquoise) and DAPI (blue) immunofluorescence staining of GFP^+^ G144 cells directly injected (injured CTX) or invaded into the dense myelin of inner cortex from the tumour bulk (invaded CTX). Dotted lines delineate the corpus callosum (CC). Scale bar = 100μm. **M**, quantification of the percentage of SOX10^+^/EdU^−^ differentiated tumour cells in the experiment shown in t. A minimum of 340 cells per xenograft were counted. Mean±SEM, n=3 xenografts per group. **p<0.01. Unpaired two-tailed Student’s t test. See also Figures S3 and S4.

Super resolution and correlative light and electron microscopy (CLEM) showed that within areas of myelin disruption tumour cells associated closely with both intact myelinated and demyelinating axons (Figure 3J, K). This did not affect tumour cell survival (Figure S3K, L), but occasionally resulted in tumour cells acquiring oligodendrocyte morphology, in line with a differentiation response (Figure S3M).

Together, this data supports a model whereby tumour infiltration into the white matter induces an injury-like microenvironment, which in turn triggers GBM differentiation in a tumoursuppressive feedback loop. To test this more directly, we carried out a time-course analysis of the response of GBM cells and the CC microenvironment to tumour infiltration. G144 xenografts were collected at 2 weeks intervals from full engraftment at 4 weeks to symptomatic disease at 12 weeks post-implanation and subjected to immunohistochemistry for differentiation and activated glia markers (Figure S4A). Tumour infiltration resulted in progressive loss of myelin integrity, which correlated with a gradual increase in microglia activation, astrocyte reactivity, OPC activation and oligodendrocyte death, all hallmarks of the glial response to brain injury (Figure S4B-L) (DiSabato et al., 2016). Importantly, tumour cell differentiation paralleled these changes, with the number of SOX10^+^/EdU^−^ tumour cells gradually increasing over time and remaining lowly proliferative, indicative of stable pre-oligodendrocyte differentiation (Figure S4L, M).

To determine whether injury is causal to tumour differentiation, we next performed gain-of-function experiments. G144 cells were injected directly into the densely myelinated inner layers of the cortex, a brain region in which tumour cells frequently infiltrate from the striatal tumour bulk, but do not differentiate (Figure 3L). Remarkably, the stab-wound injury caused by the needle (evidenced by induction of astrocyte reactivity and microglia activation, which was absent in the infiltrated cortex, Figure S4N-P) was sufficient to induce GBM cell differentiation to SOX10^+^/EdU^−^ cells (Figure 3L, M). Thus, GBM differentiation is a white matter injury-like response.

### Differentiation is white matter-dependent

To determine the stability of white-matter induced differentiation, we carried out secondary xenografts. G144 cells were isolated from CC, ST and B xenograft regions as above, and immediately re-injected in the striatum of secondary hosts (Figure 4A). Survival analysis indicated that all cells, regardless of region of origin, formed tumours with similar latency, indicative of comparable tumourigenic potential (Figure 4B). Furthermore, independent of initial SOX10 levels, all secondary lesions recapitulated SOX10 expression patterns of the primary tumours, which was high in white matter and low in grey matter regions and bulk (Figure 4C). This suggests that once removed from white matter, pre-oligodendrocyte-like tumour cells may de-differentiate back to a GSC state. To directly test this hypothesis, we used the surface marker O4 together with GFP to FACS-purify pre-oligodendrocyte tumour cells from myelin-rich CC and B regions of xenografts, seeded them acutely *in vitro* in the presence of mitogens and monitored their morphology and proliferation in real-time by time-lapse microscopy (Figure S5A, B). Consistent with the *in vivo* results, we found that approximately 20% of branched, pre-oligodendrocyte O4^+^ tumour cells retracted their processes, re-acquired GSC morphology, and re-entered the cell-cycle (Figure S5B and Videos S2 and S3). Thus, maintenance of pre-oligodendrocyte-like fate *in vivo* requires continuous exposure to white matter.

**Figure 4.**
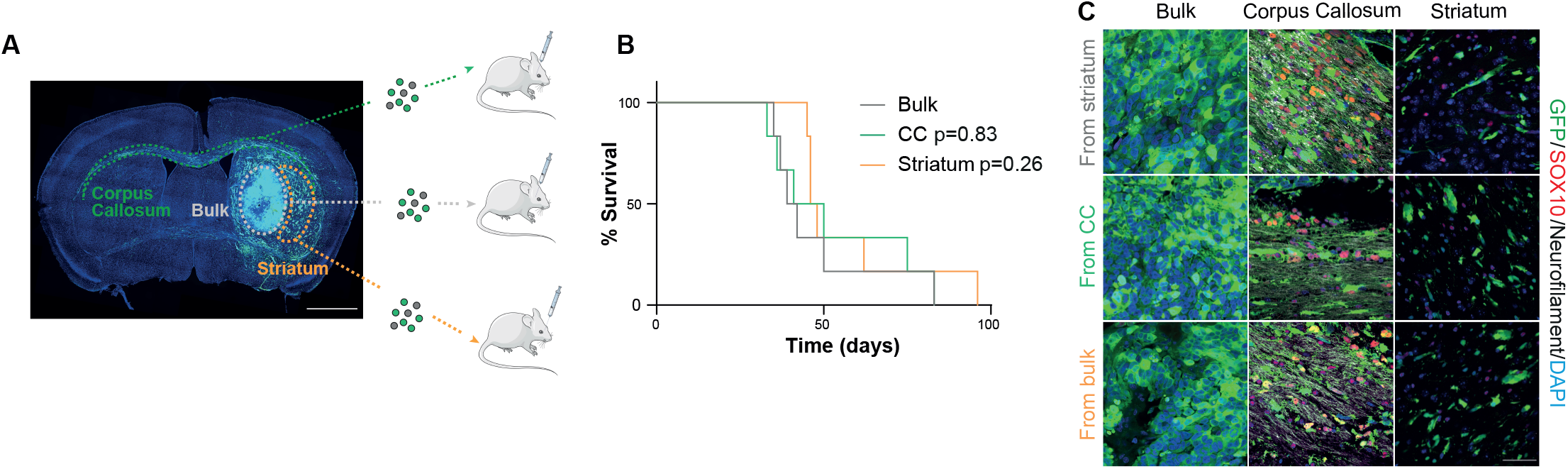
Differentiation depends on continuous exposure to white matter. **A**, schematic representation of experimental workflow. **B**, Kaplan Meier survival plot of nude mice injected with G144 cells acutely isolated from the B, CC and ST of primary xenografts. n=6 mice/group. **C**, representative images of secondary GFP^+^ tumours stained for neurofilament (grey) to identify axonal bundles, SOX10 (red) and DAPI (blue). Tumour areas from the indicated brain regions are shown revealing a differentiation pattern identical to primary lesions. Scale bar=50μm. See also Figure S5.

### SOX10 drives differentiation

In normal oligodendrogenesis, SOX10 is essential for lineage progression, differentiation and myelination (Hornig et al., 2013; Stolt et al., 2002; Turnescu et al., 2018). We therefore asked whether the increase in SOX10, which marks differentiated tumour cells in white matter may also play a causal role in the induction of tumour cell differentiation. To address this, we transduced G144 cells with an inducible Tet-ON SOX10 overexpression construct and profiled vehicle- and doxycycline (Dox)-treated cultures by RNA-seq at 48h post-induction. This time point was chosen to maximise the specificity of detected transcriptional changes and enrich for direct SOX10 targets. Immunofluorescence analysis confirmed detectable SOX10 protein expression in approximately 60% of the cells (Figure S6A, B). We identified 186 differentially expressed genes, the majority of which were upregulated (157, Table S7). Approximately 40% of SOX10-induced genes overlapped with genes upregulated in the corpus callosum *in vivo*, alongside significant overlap with signatures of oligodendrocyte lineage cells (Figure 5A). These genes included both immature and mature oligodendrocyte markers, such as GPR17, ERBB3 and the myelin genes PLP1, CLDN11, MYRF and UGT8 (Figure 5B). Furthermore, GO analysis identified enrichment for terms associated with oligodendrocyte differentiation, including axon ensheathment, myelination, cell adhesion and positive regulation of gliogenesis (Figure 5C and Table S8). Thus, SOX10 controls a transcriptional programme of oligodendrogenesis in tumour cells.

**Figure 5:**
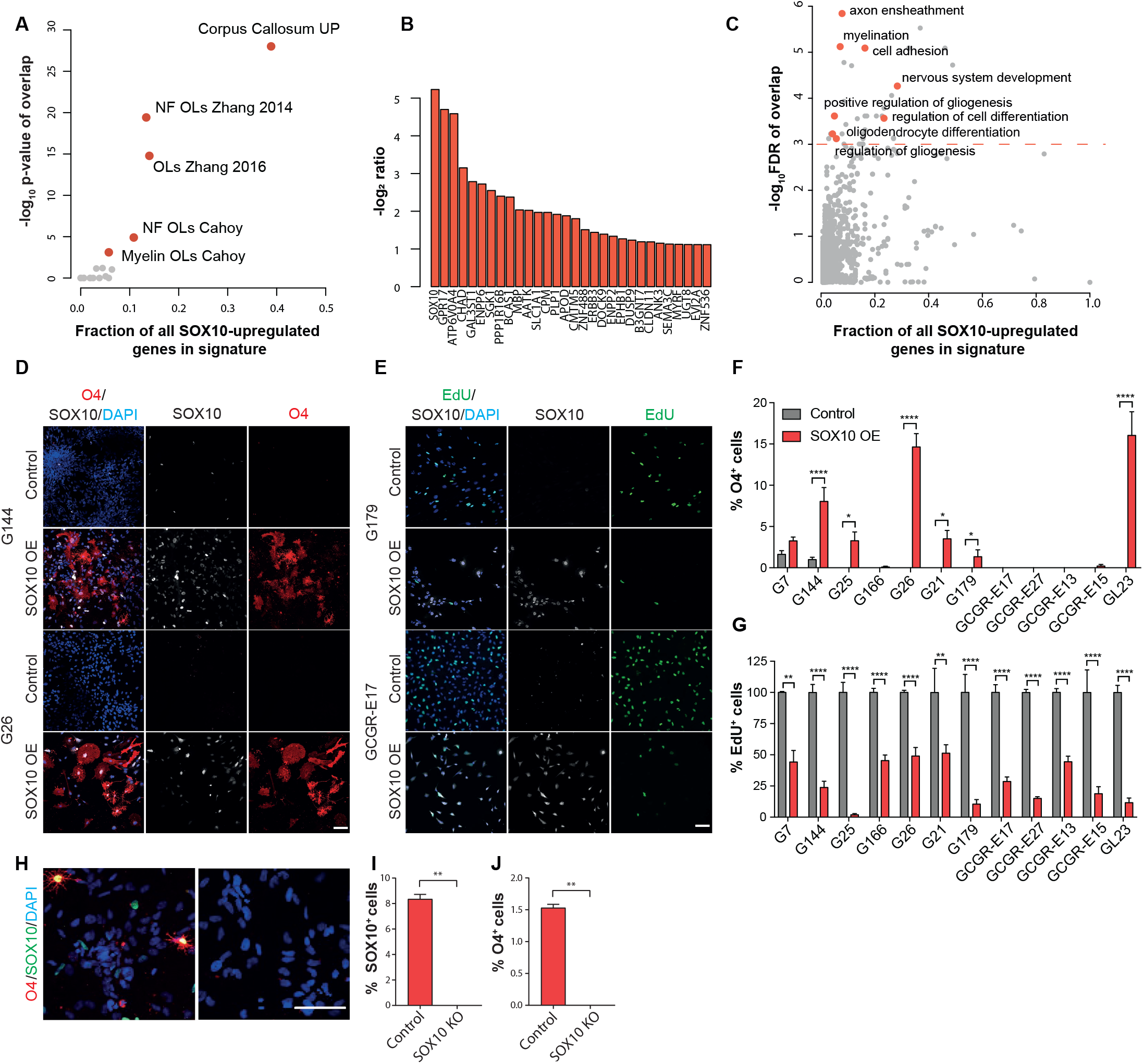
SOX10 overexpression induces pre-oligodendrocyte differentiation. **A**, Enrichment of the genes positively regulated by SOX10 *in vitro* for signatures of glial and neuronal lineages (Table S4) and a list of genes up-regulated in G144 cells upon invasion into the Corpus callosum *in vivo* (Corpus callosum UP). −log_10_ transformed p-values of one-sided Fisher tests are plotted as a function of the percentage of all SOX10 regulated genes present in each list. **B**, most up-regulated genes upon SOX10 induction *in vitro* (top 20). DEseq2 log_2_ ratio of Doxycyclin-versus vehicle-treated cells are plotted. **C**, GO enrichment analysis of the genes positively regulated by SOX10 *in vitro* (Table S8). GO terms belonging to the “Biological process” ontology are shown and selected terms are highlighted. −log10 transformed FDR q-values of overlaps are plotted as a function of the percentage of all SOX10 regulated genes present in each list. **D**, O4 (red), SOX10 (grey) and DAPI (blue) staining of indicated GSC lines transduced with empty (Control) or constitutive SOX10-encoding lentiviral vectors. Scale bar=100μm. **E**, representative images of indicated lines transduced with control or SOX10 lentivirus and stained for EdU (green), SOX10 (grey) and DAPI (blue). Scale bar=100μm. **F**, quantifications of percentage of O4^+^ pre-oligodendrocyte cells in cultures shown in d. A minimum of 200 cells across duplicate coverslips was counted per biological repeat. Mean±SEM, n= 3 independent transductions. ***p<0.001, **p<0.01, *p<0.05. Two-way ANOVA with Sidak's multiple comparisons test. **G**, quantifications of percentage of proliferating (EdU^+^) cells in the same cultures as in e. A minimum of 200 cells across duplicate coverslips was counted per biological repeat. Mean±SEM, n=3 independent transductions ***p<0.001, **p<0.01, *p<0.05. Two-way ANOVA with Sidak's multiple comparisons test. **H**, representative immunofluorescence images of Control and SOX10 knock-out G144 cultures differentiated for 14 days by growth factor withdrawal and stained for SOX10 (green) and O4 (red). **I**, **J**, quantifications of the cultures in h. A minimum of 250 cells across duplicate coverslips was counted per biological repeat. Mean±SEM, n=3 independent cultures **p<0.01, *p<0.05. unpaired two-tailed Student’s t test. See also Figure S6.

To functionally assess effects of this programme on tumour cell phenotypes, we overexpressed SOX10 in a panel of patient-derived GSC lines. This included two lines with ability to spontaneously differentiate to pre-oligodendrocyte-like cells (G7, G144). Cells were transduced with lentiviral vectors expressing constitutive SOX10 (SOX10 OE), or control empty vectors, and subjected to differentiation by growth factor withdrawal for 7 days (Conti et al., 2005). SOX10 overexpression induced 7 of the 12 lines to generate O4^+^ pre-oligodendrocytes, regardless of basal differentiation competency, and, remarkably, decreased proliferation in all lines, as measured by EdU incorporation (Figure 5D-G). This indicates that SOX10 is sufficient for GBM cell differentiation. To determine if it is also necessary, we knocked-out SOX10 by CRISPR/Cas9-mediated gene editing in G144 cells and carried out differentiation assays as above. We found that differentiation to O4^+^ cells was fully abolished in the absence of SOX10 (Figure 5H-J). Together, these experiments demonstrate that SOX10 functions as a master regulator of pre-oligodendrocyte fate in GBM and confirm that corpus callosum phenotypes *in vivo* are largely mediated by SOX10 upregulation.

### Increased SOX10 suppresses tumourigenesis

Our results so far suggest that manipulations capable of increasing SOX10 levels white matter-independently should suppress tumourigenesis by promoting stable differentiation throughout the tumour mass. To test this more directly, we performed intracranial transplantations of luciferase and GFP-tagged G144 cells transduced with SOX10 OE or Control lentiviruses and monitored tumour growth and disease-free survival. SOX10 overexpression significantly delayed tumourigenesis, resulting in an overall increase in median survival from 65 to 91 days (Figure 6A, B). To understand the mechanisms responsible, we examined Control and SOX10 OE tumours at 4 weeks post implantation by immunofluorescence analysis. EdU labelling revealed a marked decrease in the total number of proliferating tumour cells in SOX10 OE compared to Control lesions (Figure 6C, D). Importantly, time-course analysis of SOX10 levels in the tumours that eventually formed revealed that SOX10 overexpressing cells were progressively outcompeted by cells that had escaped transduction, confirming that high SOX10 levels are tumoursuppressive (Figure S7A).

**Figure 6:**
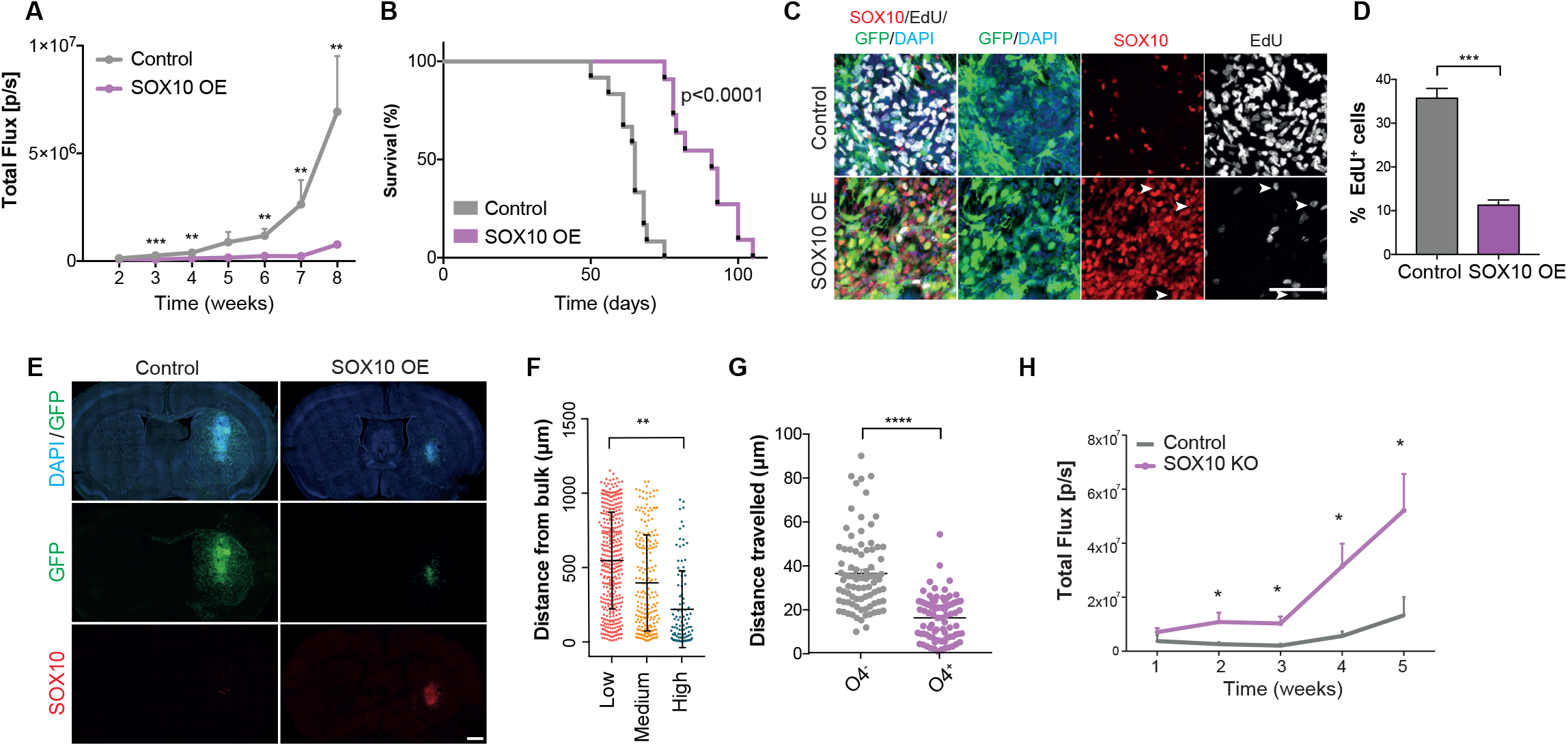
SOX10 overexpression delays tumourigenesis. **A,** quantification of luciferase bioluminescence radiance measured at the indicated time points in both groups n=12 for Control and 13 for SOX10 OE, error bars denote SEM. Unpaired two-tailed Student’s t test. **B**, Kaplan Meier survival plot of mice injected with GFP^+^ G144 Control and SOX10 OE cells n=8 for Control and 10 for SOX10 OE. **C**, SOX10 (red), EdU (grey) and DAPI (blue) immunofluorescence staining of GFP^+^ Control and SOX10 OE tumours at 4 weeks post-injection. Arrowheads denote EdU^+^ cells that are negative for SOX10. **D**, quantification of percentage of EdU^+^/GFP^+^ tumour cells in Control and SOX10 OE lesions from C. A minimum of 200 cells was quantified across 2 ROIs per mouse. Mean±SEM are plotted. n=3 mice/group ****p<0.0001. Unpaired Student’s t test. **E**, representative images of GFP^+^ G144 Control and SOX10 OE tumours stained for SOX10 (red) and DAPI (blue) at 4 weeks post-injection. **F**, quantification of migrated distance from tumour bulk edge of tumour cells expressing low, medium or high SOX10 levels in SOX10 OE tumours at 4 weeks post-injection. A minimum of 200 cells was quantified per mouse. Each dot represents a cell. Mean±SEM are indicated. n=3 mice/group. One-way ANOVA. **G**, migrated distance of O4^+^ and O4^−^ tumour cells acutely isolated from the corpus callosum region of G144 xenografts and cultured for 72h in neural stem cell conditions. Each dot represents a cell, n=90 cells/group pooled from 3 independent experiments. Median±SEM are shown. n=3 tumours *p<0.05. Unpaired two-tailed Student’s t test. **H**, Quantification of luciferase bioluminescence radiance of Control and SOX10 knock-out (SOX10 KO) G144 patient-derived xenografts measured at the indicated time points. n=5 Control and n=7 SOX10 KO tumours. Error bars denote SEM. Unpaired two-tailed Student’s t test. See also Figure S7.

Although SOX10-mediated differentiation was most pronounced in invasive tumour cells that infiltrate the corpus callosum (Figure1C-E), normal oligodendrocyte maturation is accompanied by a progressive loss of migratory potential (Barateiro and Fernandes, 2014; de Castro et al., 2013). We therefore sought to understand the impact of differentiation on GBM invasion by imaging whole sections of 4 weeks tumours. Surprisingly, SOX10 OE tumours appeared less diffuse and migrated much shorter distances than controls (Figure 6E and Figure S7B). In addition, within SOX10 OE tumours the majority of invasive tumour cells had low or undetactable SOX10 levels, whereas SOX10 high cells were found predominantly closest to the tumour bulk (Figure 6F). These results indicate that differentiation also reduces the migration of GBM cells. To determine whether this phenotype was cell-intrinsic, we measured *in vitro* motility of SOX10 OE and Control G144 cells by live cell imaging and found that increased SOX10 levels profoundly reduced cell motility (Figure S7C-E). This was not an artefact of SOX10 overexpression, as O4^+^ tumour cells acutely FACS-purified from the corpus callosum of wildtype G144 xenografts were also significantly less motile than O4^−^ cells from the same region (Figure 6G and Figure S7F, G).

These results suggest that white matter effects are tumour suppressive and might slow down the progression of the primary disease. To test this hypothesis experimentally, we compared the tumourigenicity of parental and SOX10 knock-out G144 cells, in which differentiation along the oligodendrocyte lineage is abolished (Figure 5H-J). We found that the growth rate of G144 cells was significantly increased upon SOX10 deletion, with knock-out cells producing larger tumours than differentiation-competent controls, as measured by bioluminescence imaging (Figure 6H). These experiments demonstrate that white matter-driven pre-oligodendrocyte maturation suppresses tumourigenesis by inhibiting proliferation and invasion of GBM cells and that in the absence of this response GBMs are more aggressive.

Finally, we explored the translational potential of these findings by testing whether known myelination-inducing pharmacological agents could differentiate GBM cells by increasing endogenous SOX10 levels. We first treated cultured G144 cells with two compounds: the clinically-approved anti-asthma medication Pranlukast and the cell permeable cAMP analog dibutyril cAMP (db-cAMP). In addition to blocking cysteinyl-leukotriene 1 receptor, Pranlukast inhibits GPR17, a negative regulator of oligodendrocyte development (Chen et al., 2009; Hennen et al., 2013; Simon et al., 2016). GPR17 is normally expressed in immature oligodendrocytes (Chen et al., 2009) and, importantly, was strongly induced in G144 cells purified from the CC and upon SOX10 overexpression (Figure 1D and 5B). db-cAMP treatment of normal progenitors results in elevated intracellular cAMP levels, which promote oligodendrocyte differentiation, likely by phenocopying GPCR activity (Mogha et al., 2016; Raible and McMorris, 1990). Treatment of G144 with either compound for 2 weeks was sufficient to increase endogenous SOX10 levels and differentiation to O4^+^ pre-oligodendrocyte cells, leading to a reduction in proliferation (Figure 7A-H). These effects were fully dependent on SOX10, as no differentiation was observed in drug-treated SOX10 knock-out cells (Figure S8A-D). Next, we administered Pranlukast to tumour-bearing mice *in vivo*. As Pranlukast has poor BBB penetration (Tang et al., 2014), we delivered it intrathecally using osmotic mini-pumps. Strikingly, we found a significant increase in the number of SOX10^+^ cells in Pranlukast-treated tumours, which was accompanied by a dramatic reduction in proliferation relative to saline-treated controls (Figure 7I-L). Pranlukast did not affect endogenous inflammatory glia, indicative of a direct effect on the tumour cells (Figure S8E-G). We conclude that a subset of GBMs can be induced to undergo pre-oligodendrocyte-like differentiation with small molecules that raise SOX10 levels and propose that such differentiation therapy may be an effective strategy for curtailing tumourigenesis and recurrence.

**Figure 7.**
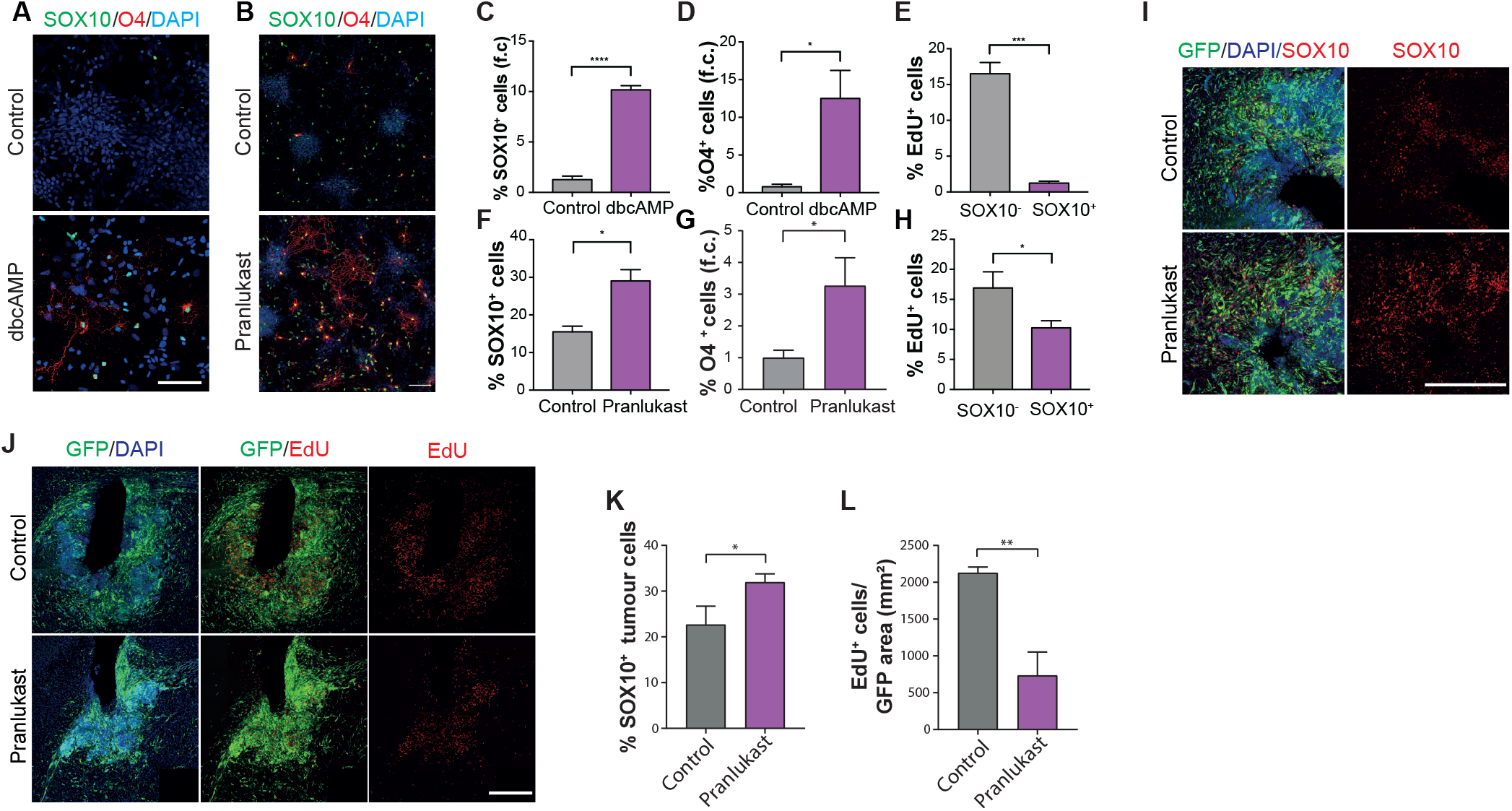
Myelination-promoting compounds suppress tumour growth. **A,** representative images of untreated (Control) and dbcAMP-treated G144 cultures stained for SOX10 (green), O4 (red) and DAPI (blue). Scale bar=100μm. **B**, representative images of untreated (Control) and Pranlukast-treated G144 cultures stained for SOX10 (green), O4 (red) and DAPI (blue). Scale bar=50μm. **C-E**, quantification of cultures in A, showing percentages of indicated populations before and after treatment. Fold change relative to control cultures (f.c.) is shown in C and D. A minimum of 1000 cells cells across duplicate coverslips was counted per biological repeat. Mean±SEM, n=3 independent cultures *p<0.05. Unpaired two-tailed Student’s t test. **F-H**, quantification of cultures in B, showing percentages of indicated populations before and after treatment. Fold change relative to control cultures (f.c.) is shown in G. A minimum of 800 cells across duplicate coverslips were counted per biological repeat. Mean±SEM, n=3 independent cultures. ***p<0.001, *p<0.05. Unpaired two-tailed Student’s t test. **I**, representative immunofluoresce images of DMSO-(Control) and Pranlukast-treated GFP^+^ G144 xenografts stained for SOX10 (red) and **J**, EdU (red). Scale bar=500μm. **K**, quantifications of number of SOX10^+^ and **L**, EdU^+^ tumour cells in the xenografts shown in I and J. Scale bar=500μm. Mean±SEM, n=3 xenografts **p<0.01. Unpaired two-tailed Student’s t test.n=3 xenografts per group. ***=p<0.01, *p<0.05. Unpaired two-tailed Student’s t test. See also Figure S8.

## Discussion

The failure of both conventional and targeted therapies is commonly attributed to the pervasive molecular intra- and intertumoural heterogeneity of GBM, which occurs at the genetic, epigenetic, transcriptional and functional levels (Brennan et al., 2013; Cancer Genome Atlas Research, 2008; Patel et al., 2014; Sottoriva et al., 2013; Sturm et al., 2012; Verhaak et al., 2010; von Neubeck et al., 2015; Wang et al., 2017). Despite this staggering diversity however, recent studies are beginning to reveal that all GBMs converge on a finite number of cellular states, which recapitulate normal developmental programmes (Lan et al., 2017; Neftel et al., 2019). Such convergence offers hope of identifying shared biological vulnerabilities that could be exploited for the treatment of many patients, independent of genetic diversity. Here we identified one such vulnerability: the competence of GBM cells to undergo pre-oligodendrocyte differentiation.

Although GSCs are known to retain the ability to partially differentiate, how differentiation is controlled in tumours has remained unclear (Caren et al., 2015; Lan et al., 2017; Neftel et al., 2019; Park et al., 2017). Our results demonstrate that differentiation occurs in specific microenvironments within the brain and identify the white matter as one such pro-differentiative niche for tumour cells with oligodendrocyte competency (Elbaz and Popko, 2019). Thus, despite the presence of extensive genetic abnormalities, exposure to appropriate environmental cues is sufficient to revert GBM cells to a more normal, differentiated phenotype, underscoring the dominance of the microenvironment in suppressing malignancy (Bissell and Hines, 2011; Weaver et al., 1997).

We found that differentiation is an injury-like response, which results from infiltrating tumour cells disrupting the white matter. Thus, although invasion along white matter is a common mode of infiltration, the ensuing myelin damage creates a tumour-suppressive feedback loop, which paradoxically slows GBM proliferation and spread (Cuddapah et al., 2014). Indeed, we showed that SOX10 knock-out GBM cells that lack oligodenderocyte differentiation competency are more tumourigenic than parental cells. This suggests that in the absence of white matter-induced differentiation, GBM would be an even more aggressive disease. Interestingly, the microenvironment of tumour-infiltrated white matter appeared remarkably similar to that of neuroinflammatory disease states, entailing severe demyelination and axonal pathology (DiSabato et al., 2016). Our EM analysis suggests that in the tumour context demyelination may be caused by both oligodendrocyte and neuron death. The underlying mechanisms remain to be determined, but a combination of compression injury to the axons, microglia and astrocytes activation, as well as excitotoxins are all likely to contribute (DiSabato et al., 2016; Seano et al., 2019).

The finding of OPC activation in tumour-infiltrated white matter suggests that the ensuing demyelinating microenvironment initiates a repair programme in normal glia. It is therefore tempting to speculate that GBM cells may recapitulate a similar oligodendrogenic programme, which results in their partial differentiation to pre-oligodendrocyte-like cells. While it is clear that an injury-like microenvironment drives differentiation, the signals and cell types responsible remain undefined. These are likely to be complex and combinatorial, but our results suggest that microglia, a key regulator of normal re-myelination, may play an important role (DiSabato et al., 2016; Lloyd and Miron, 2019). Indeed, we found that microglia activation was selective to tumour white matter and immediately preceded tumour cell differentiation. It would be of great interest to examine the contribution of microglia to GBM differentiation in future studies.

Our findings are of clinical relevance as we show that exploiting the aberrant GBM injury-like response through white matter-independent upregulation of SOX10, locked tumour cells in the differentiated, non-proliferative state and suppressed tumourigenesis. Furthermore, our work predicts that myelination-promoting compounds, should be particularly effective in tumours with oligodendrocyte lineage competency, providing a potential strategy for patient stratification. Future studies should continue to explore how the intersection of repair and developmental programmes may afford novel translational opportunities for GBM.

## Acknowledgements

This work was funded by Cancer Research UK (A21203; S.P., L.B., L.Z., D.K.) and the Medical Research Council (S.P., S.M.), the Medical Research Council funding to the MRC LMCB University Unit at UCL, MC_U12266B (J.J.B.). It used the computing resources of the UK Medical Bioinformatics partnership (UK MED-BIO; aggregation, integration, visualization, and analysis of large, complex data), which is supported by the Medical Research Council and Imperial College High Performance Computing Service. The Glioma Cellular Genetics Resource is funded by a Cancer Research UK Accelerator Award (A21992). We thank W. Richardson and H. Li for reagents, F. Coutinho for histology, J. Manji for help with microscopy, T. Adejumo and J. Elliott for help with FACS, S. Khadayate for help with bioinformatics and L. Game for technical advice.

## Author Contributions

L.B. and S.P. designed, conducted, and analysed experiments. M.C., D.K, L.Z., L.R., S.C.D., M.W. and M.E. conducted and analysed experiments. J.J.B. acquired and analysed electron microscopy data. Z.J. and S.B. provided patient samples and contributed to immunohistological analysis. S.M.P. and G.M provided patient-derived lines. P.B.D. provided xenograft samples. S.M. conducted bioinformatics analysis and helped with study design. L.B. and S.P. wrote the manuscript. S.P. conceived the project and supervised all aspects of the work.

## Methods

### Animals

All procedures were performed in compliance with the Animal Scientific Procedures Act, 1986 and approved by with the UCL Animal Welfare and Ethical Review Body (AWERB) in accordance with the International guidelines of the Home Office (UK). C57Bl6 and immunocompromised mouse lines were purchased from Charles River.

### Patient-derived xenograft models

Xenografts were performed using CD-1 nude mice for G144 and NOD-SCID-IL2R gamma chain-deficient (NSG) for all other lines. For all tumour studies 8-12 week old male or female immunocompromised mice underwent stereotactic implantation of GSC lines using 1×10^5^ cells with the exception of G144 reinjection experiments for which 5x×10^4^ were used (anteroposterior 0, mediolateral −2.5, dorsoventral −3.5; corpus callosum: anteroposterior 0, mediolateral −1, dorsoventral −1.75; inner cortex: anteroposterior 0, mediolateral −1, dorsoventral −1; outer cortex: anteroposterior 0, mediolateral −1, dorsoventral −0.5). Tumour growth was monitored using an IVIS Spectrum *in vivo* imaging system (Perkin Elmer). 10 minutes following i.p. D-luciferin (120 mg/kg, Intrace medical) bioluminescent images were acquired under isoflurane anaesthesia. Tumour size was quantified by calculating total flux (photons/s/cm^2^) using Living Image software (Xenogen, Caliper Life Sciences). Animals were sacrificed and tumours collected when they showed signs of distress or >10% weight loss. For experiments shown in Figure 4 c-f and r-u tumours were collected at 4 weeks post-implantation. To assess tumour cell proliferation EdU was administered by i.p. injection (50mg/kg) 4 hours prior to collection. Survival was analysed using the Kaplan-Meier method and significance calculated using the log-rank Mantel-Cox test.

### Cell Culture

Patient-derived GCGR cell lines were provided by the glioma cellular genetics resource: (www.gcgr.org.uk). All GSC lines were cultured adherently in serum-free GSC media (N2 (1/200), B27 (1/100) (Life Technologies), 1 mg/ml laminin (Sigma), 10 ng/mL EGF and FGF-2 (Peprotech), 1x MEM NEAA (gibco), 0.1 mM betamercaptoethanol, 0.012% BSA (Gibco), 0.2 g/L glucose (Sigma), 1000 U/ml penicillin-streptomycin (Sigma)) as previously described(Pollard et al., 2009). Medium was changed every 3 d, and cells were dissociated using Accutase solution (Sigma). For induction of SOX10 expression, doxycycline was used at a concentration of 3μg/ml. For experiments involving assessment of proliferation, cells were exposed to 10μm EdU for 4 hours prior to collection.

### FACS of tumour cells

Brains were harvested and tumour regions corresponding to tumour bulk (B), invaded striatum (ST) and corpus callosum (CC) were micro-dissected under fluorescent guidance in ice cold HEPES buffered HBSS. Micro-dissected regions were dissociated by incubating with papain (20 units/ml) DNase (0.005%) for 30min at 37°C (Worthington). Samples were triturated in HEPES buffered EBSS to fully dissociate the tissue and centrifuged (3 min, 300xg). Digestion was terminated by trituration in EBSS ovomucoid inhibitor (10mg/ml), albumin (10mg/ml) and DNase (0.005%). Following centrifugation (3 min, 300xg), cells were resuspended in 400μL FACS buffer (1.5% BSA, 2.5mM HEPES, 1mM EDTA in PBS) containing 2.5% RNAsin and 1/10000 DAPI (Promega). Cells for re-injection experiments were isolated as above and subjected to a further two centrifugation steps (1min at 200xg) in 10ml warm EBSS after digestion to reduce myelin debris. For isolation of O4^+^ tumour cells, cells were dissociated as described above. Following centrifugation in EBBS, cells were resuspended in 500ul GSC media containing 1:500 mouse anti-O4 (Alexa Fluor 594) and incubated for 15min at 37°C. Cells were washed once in PBS + 3% BSA and resuspended in 400μL FACS buffer containing 1/10000 DAPI. O4^+^ and O4^−^ populations were seeded in serum-free GSC media containing EGF and FGF for live cell imaging.

### Whole-transcriptome amplification, library construction, sequencing and processing

For RNA-seq of FACS-sorted tumour cells, dsDNA libraries were prepared according to the Smart-seq2 protocol from FACS-sorted GFP+ cells (RIN >8)(Picelli et al., 2014). Next-generation sequencing libraries were prepared from 1ng dsDNA using the Nextera XT DNA library preparation kit (Illumina) and indexed using the Nextera XT index Kit (Illumina). For RNA-seq of *in vitro* GSCs, oligo dT-based mRNA isolation was performed using the NEBNext Poly(A) mRNA magnetic isolation module. dsDNA libraries were prepared using NEB Ultra II directional RNA library prep kit and indexed using NEBNext^®^ Multiplex Oligos. All libraries were diluted to a final concentration of 2.5nM, pooled and sequenced on an Illumina HiSeq 2500 instrument. Raw data were processed using RTA version 1.18.64, with default filter and quality settings. The reads were demultiplexed with CASAVA 1.8.4 (allowing 0 mismatches). Raw reads were aligned to the human genomes (hg38) and assigned to genomic features both using the STAR aligner(Dobin et al., 2013). After filtering, only genes with at least 5 counts in at least 3 samples were included in the final dataset (n=16044 for in vivo RNA-seq, n=14621 for in vitro RNA-seq).

### Analysis of RNA-seq data

Differential expression analysis was performed and normalized counts were generated using the DESeq2 Bioconductor package (Table S1) (Love et al., 2014). For Figure 1c, genes significantly regulated (P_adjust_ < 0.1) and with an absolute DESeq2 log2 ratio >0.58 in at least one of the Corpus callosum vs Bulk or Striatum vs Bulk comparisons were selected and clustered using K-means (7 clusters). Overlap of each cluster with gene signatures specific for proliferating cells and a series brain cell types was assessed using a one-sided Fisher exact test and p-values were corrected for multiple testing using the Benjamini-Hochberg approach. For Figure S1F, genes belonging to each gene signatures and significantly regulated (p_adjust_ < 0.1) in each comparison are shown. Brain signatures were generated using one human and two mouse transcriptomics datasets(Cahoy et al., 2008; Zhang et al., 2014; Zhang et al., 2016). Genes with multiple entries were averaged and low coverage genes filtered out. Condition replicates were averaged, data median centred before the most variable genes between cell types were selected (Coefficient of variation, CV >1 for Zhang 2016, CV >1.5 for Zhang 2014, CV > 0.1 for Cahoy 2008) for K-means clustering (Figure S1A-C). The cut-offs on coefficient of variation and the numbers of K-means clusters were selected empirically to maximise the discrimination between cell types. We defined single clusters with good cell type discrimination as gene signatures (Figure S1 A-C, Table S3). Proliferating cells signatures were obtained from the GSEA website. Gene expression z-scores of the SOX10 mRNA were downloaded from cBioportal for samples from the TGCA cohort. GBM subtypes for each sample were retrieved from Wang *et al*, 2017^4^. For Figure 3c, GO enrichment analysis was performed using the VLAD on-line portal (http://proto.informatics.jax.org/prototypes/vlad/).

### Data availability

RNA-seq data that support the findings of this study have been deposited in GEO (accession codes pending). The authors declare that all other data supporting the findings of this study are available within the paper and its supplementary information files.

### SOX10 overexpression

Constitutive SOX10 expression for analysis of differentiation and proliferation *in vitro* as well as *in vivo* tumorigenesis studies was achieved by gateway cloning of human SOX10 from pDONR221-SOX10 into a pLenti-CMV-BLAST-DEST vector. For the empty control vector, the ccdB gene was removed using EcoRV and HpaI blunt end ligation. For *in vitro* RNA-seq an inducible SOX10 construct was generated using gateway cloning to insert SOX10 from pDONR221-SOX10 into pCW57.1. pDONR221-hSOX10 was a gift from William Pavan (Addgene plasmid #24749; http://n2t.net/addgene:24749; RRID:Addgene_24749)(Cronin et al., 2009). pCW57.1 was a gift from David Root (Addgene plasmid #41393; http://n2t.net/addgene:41393; RRID:Addgene_41393). Lentivirus was produced by cotransfecting with the HIV-1 packaging vector Delta8.9 and the VSVG envelope glycoprotein into 293T cells using *polyethylenimine*. Virus was concentrated by ultracentrifugation (3 hr, 50,000xg, 4°C).

### Cell Migration assay

Live cell imaging was performed using either the Zeiss Live Cell Imager Z1 or the IncuCyte Zoom. Cells were tracked manually in ImageJ using the ManualTracking plugin, migration distance was calculated and cell trajectories were visualized using the Chemotaxis and Migration Tool provided by Ibidi.

### Quantitative RT-PCR

RNA was extracted using Trizol Reagent (Sigma). RNA was reverse transcribed using iScript gDNA clear cDNA synthesis kit (Bio-rad) and quantitative PCR was performed using the qPCRBIO SyGreen Mix Lo-Rox (PCR Biosystems). Relative expression values for each gene of interest were obtained by normalizing to GAPDH. Primers used were Fw 5’-CCTCACAGATCGCCTACACC-’3 and Rev 5’-CATATAGGAGAAGGCCGAGTAGA-’3 for SOX10 and Fw 5’-GTCTCCTCTGACTTCAACAGCG-’3 and Rev 5’-ACCACCCTGTTGCTGTAGCCAA-’3 for GAPDH.

### Immunofluorescence and Immunohistochemistry

For immunohistochemical analysis, mice were perfused with 4% PFA and the brain post-fixed in 4% paraformaldehyde overnight at 4°C. 50μm vibratome sections were permeabilised and blocked in 1% Triton X-100, 10% serum for 1.5hrs then incubated overnight at 4°C with primary antibodies diluted in 0.1% Triton X-100, 10% serum. For paraffin embedded sections, following deparaffinisation, antigen retrieval was performed using in citrate buffer (Sigma) and sections were permeabilised and blocked 1% Triton X-100, 10% serum for 1hr then incubated overnight at 4°C with primary antibodies diluted in 0.1% Triton X-100, 10% serum.

For *in vitro* immunofluorescence, cells were fixed in 4% PFA, permeabilised in 0.5% Triton X-100 and blocked in 10% serum in PBS then incubated with primary antibody overnight at 4°C in PBS + 10% serum. Anti-O4 antibody staining was performed on live cells for 30min at 37°C in cell culture media. Secondary antibodies were diluted in 10% serum in DAPI (1:10000 in PBS) and incubated at RT for 1h. Analysis of EdU incorporation was performed using Invitrogen’s™ Click/iT™ EdU imaging Kit according to the manufacturer’s specifications. Imaging was carried out using the Zeiss Z1 upright microscope or the Zeiss LSM880 confocal microscope. Quantification was performed using Fiji ImageJ.

Primary antibodies used were mouse anti-CC1 (1:1000, abcam ab16794), rabbit anti-Ki67 (1:250, abcam ab16667), goat anti-MBP (1:1000, santa cruz sc-13912), rat anti-MBP (1:500 paraffin; 1:1000 coverslips, sigma MAB386), chicken anti-neurofilament (1:2000, abcam ab4680), mouse anti-HuNu (1:250, sigma MAB1281), mouse anti-O4 (1:500, R&D MAB1326), rabbit anti-RFP (1:500, antibodies online AA234 (ABIN129578)), rabbit anti-OSP (1:500, abcam ab53041), goat anti-SOX10 (1:1000, R&D AF2864), rat anti-CD68 (1:500, abcam ab53444), rabbit anti-Iba1 (1:1000, Wako 019-19741), mouse anti-Sox2 (1:100, abcam ab79351), rabbit anti-Sox2 (1:1000, abcam ab97959), rat anti-CD44 (IM7) (1:500, Invitrogen 14-0441-81), rabbit anti-GFAP (1:1000, Dako Z0334), mouse anti-CNP (1:500 abcam ab6319). Alexa Fluor conjugated secondary antibodies were obtained from Thermo Fisher.

### Image processing and quantifications

All image quantifications were carried out using Fiji ImageJ. For analysis of SOX10 expression in infiltrating G144 tumour cells, white matter (corpus callosum) and grey matter regions containing at least 750 NuMa^+^cells were defined for quantification across n=3 mice and SOX10^+^ and SOX10^−^ human cells counted within these regions. For analysis of EdU incorporation a minimum of 500 NuMA^+^ cells was quantified for SOX10 expression and EdU positivity across 2 independent ROIs of infiltrated corpus callosum. For analysis of SOX10^+^/ EdU^−^ cells upon implantation into the white matter of the corpus callosum (n=3) or grey matter of the upper cortex (n=3) a mimumum of 140 cells per xenograft were counted. For analysis of regional EdU^−^ incorporation in GFP^+^ tumour cells of G144 xenografts (n=3), a minimum of 200 cells were counted per xenograft. For analysis of the impact of PDGFRA signalling on oligodendroglial differentiation *in vitro*, a minimum of 3500 cells were counted per biological repeat (n=4). For quantification of the percentage of SOX10^+^/HuNu^+^ tumour cells in white and grey matter of all other patient-derived xenografts n=4, a minimum of 550 HuNu^+^ cells was quantified across 2 independent ROIs within white matter (corpus callosum) or grey matter. Analysis of tumour cell proliferation within the corpus callosum of xenografts PDX 23, 67, 523 and 800 was carried out for a minimum of 500 SOX10^+^ or SOX10^−^ HuNu^+^ cells across 2 independent ROIs. For analysis of SOX2^+^ tumour cells undergoing proliferation (Ki67^+^) in patient tissue, a minimum of 300 cells were quantified across 2 independent ROIs for each case n=3.

For quantification of percentage of SOX10^+^ tumour cells in areas of high and low myelin disruption, a minimum of 120 cells was quantified across a minimum of 6 ROIs selected within intact or disrupted white matter for n=4 xenografts. Myelin disruption was defined as white matter regions containing Neurofilament^+^ axons and Fluromyelin staining intensity <25% of contralateral intact regions. For analysis of endogenous oligodendrocytes (SOX10^+^/NuMA^−^) within the corpus callosum as a function of number of invaded SOX10^+^ G144 (SOX10^+^/NuMA^+^) tumour cells, quantification was carried out across a minimum of 2 ROIs and counting >1000 cells per mouse for n=5 tumours and n=2 intact control brains.

For quantification of the percentage of GFP^+^ tumour cells in contact with axons within these regions a minimum 300 SOX10^+^ or SOX10^−^ cells was quantified across 3 independent ROIs for n=3 tumours. Contact was defined as <1.5μm from the edge of the nucleus to a Neurofilament^+^ axon. For analysis of microglia within the white matter (corpus callosum), grey matter (cortex) or tumour bulk a minimum of 140 cells per region per xenograft were counted. Analysis of endogenous oligodendrocytes was conducted across 5 xenografts and 2 control brains. A minimum of 450 cells were counted across 2 ROIs/xenograft. Quantifications of cleaved caspase-3 staining of GFP^+^ G144 tumour cells within the white matter (corpus callosum), grey matter (cortex) or tumour bulk was conduceted for a minimum of 200 cells per region per xenograft (n=3).

For timecourse analysis of corpus callosum fluromyelin intensity, mean gray values were normalised to the max gray values to account for variation in image intentity. For all timecourse analyses a minimum area of 300μm^2^ was analysed for a minimum of 3 xenografts. For analysis of SOX10^+^ EdU^−^ cells within the invaded of injured cortex a minimum of 340 cells were counted per xenograft (n=3-4). For analysis of microglia within the invaded or injured cortex a minimum of 90 cells were counted. For quantification of the percentage of GFP^+^ tumour cells in contact with axons within these regions a minimum 300 SOX10^+^ or SOX10^−^ cells was quantified across 3 independent ROIs for n=3 tumours. Contact was defined as <1.5μm from the edge of the nucleus to a Neurofilament^+^ axon. For quantifications of the percentage of O4^+^ pre-oligodendrocyte cells and EdU^+^ proliferating cells in control and SOX10 transduced cultures of GSC lines a minimum of 200 cells across duplicate coverslips were counted per biological repeat for n=6 independent cultures per line. For analysis of differentiation in SOX10 knock-out cells a minimum of 250 cells were counted across duplicate coverslips per repeat. For SOX10 induction in vitro a minimum of 90 cells per group across 2 independent cultures on triplicate coverslips. For quantifications of the relative proportions of SOX10^−^, low or high tumour cells at different timepoints (pre-implantation, 4 weeks and survival (11-12 weeks) a minimum of 270 cells were analysed. SOX10 status was assessed used mean gray values of individual nuclei. SOX10-cells were counted as those with a mean gray value less than background. SOX10 high threshold was determined by measuring the minimum mean gray value of >20 cells showing high SOX10 expression.

Proliferation of control or SOX10 OE tumours was quantified across n=3 tumours at 4 weeks post-implantation, counting a minimum of 200 cells from 2 ROIs per mouse. Quantification of invasion of n=3 tumours was performed by measuring the total area occupied by tumour cells and normalising it to the perimeter of the tumour bulk to account for different rates of tumour growth. Invasion was further analysed within SOX10 OE by measuring the distance of SOX10 low, medium and high expressing cells from the tumour bulk. SOX10 low and high thresholds intensities were defined as the 25^th^ and 75^th^ percentiles respectively based on analysis of the average intensities across >100 nuclei within the tumour bulk. A minimum of 200 cells was quantified per mouse. For *in vitro* quantifications of the percentage of O4^+^ and EdU^+^ cells following exposure to pranlukast or control, a minimum of 800 cells across duplicate coverslips were counted for n=3 independent cultures. For dbcAMP experiments a minimum of 900 cells was counted across duplicate coverslips for n=3 independent cultures. For in vivo quantifications of the percentage of SOX10^+^ GFP^+^ tumour cells in control of pranlukast treated tumours a minimum of 1200 cells were counted. For analysis of proliferation total EdU^+^ nuclei were normalised to the total GFP+ tumour area (n=3). For analysis of microglia a minimum of 420 cells were counted across n=3 xenografts.

### Targeted electron microscopy

GFP-labelled G144 xenografts were perfusion fixed with 4% formaldehyde, and then further immersion fixed for 8 hours. Vibrating microtome sections (100μm) were immunolabelled for Neurofilament and SOX10 and imaged by confocal to map the location of the tumour within the section. For correlative light and electron microscopy (CLEM) of specific SOX10 positive GFP labelled cells, vibratome sections were mapped using the 20X objective to identify regions of interest. A small asymmetric piece of tissue (<1mm) containing the region of interest was further dissected from the Vibratome section and was completely mapped using the 63X objective. Vibratome sections were then processed for electron microscopy essentially as described for peripheral nerve tissue(Cattin et al., 2015). Briefly, sections were further treated sequentially with formaldehyde: glutaraldehyde, osmium tetroxide: potassium ferricyanide, osmium tetroxide, thiocarbohydrazide, uranyl acetate and lead aspartate prior to dehydration through an ethanol series and embedding in Epoxy resin. For CLEM samples, care was taken to ensure that the piece of tissue was mounted flat, in the correct orientation, to guarantee targeting the cells of interest imaged by confocal microscopy. Serial ultrathin sections (70nm) were taken using a diamond knife (Diatome) and an ultramicrotome (UC7, Leica) and collected on formvar coated slot grids. Sections were imaged in a transmission electron microscope (T12 BioTwin, Thermofisher) and captured with a CCD camera running iTEM software (Morada, Olympus SIS). All EM analysis was conducted on a minimum of 50 axons (n=3-4). Axons were considered to have decompacted myelin when >15% of the axonal circumference exhibited decompaction of the associated myelin. Degenerating axons were scored as those exhibiting any of the following features: condensed axoplasm, organelle accumulation, axonal swelling, vacuoles, dark axoplasm. G ratios were calculated by dividing the axonal diameter by the corresponding axonal + myelin sheath diameter. Feret diameters were used to account for the imperfect circularity of axons.

### Neuropathological assessment of Sox10 expression in human GBM

SOX10 expression pattern was investigated in brain tissue samples of three glioblastoma patients operated at the National Hospital for Neurology and Neurosurgery, UCL Hospitals Foundation Trust between 2009 and 2017. None of the patients had radiotherapy or chemotherapy prior to surgery, and they did not have any relevant comorbidities. Patients consent was obtained for the use of all samples. The project received ethical approval from London - Queen Square Research Ethics Committee (08-077). All 3 tumours were wildtype for IDH1 and IDH2 mutations and were the original lesions from which lines GCGR-L12, GL67 and GL23 were isolated. Clinical information is provided in Table S6.

For each case the resection material was reviewed and the region containing tumour infiltration in the surrounding white matter was selected for the analysis. The resected tissues were immediately fixed in 10% buffered formalin and processed into paraffin blocks using standard methods in the Division of Neuropathology, NHNN.

### Statistics

Statistical analysis was performed using GraphPad Prism 7.0. All data are expressed as mean ± SEM. Significance was calculated using 1- or 2-tailed Student’s *t* test, ANOVA with Bonferroni post-hoc test, Two-way ANOVA with Sidak's multiple comparisons tests or Pearson’s correlation as indicated in the figure legends. No statistical method was used to predetermine sample size. Sample size was determined based on existing literature and our previous experience. Shapiro-Wilk test was used to confirm normal distribution of the data.

