## Supplemental information for "The white matter is a pro-differentiative microenvironment for glioblastoma"

### Supplemental Figure Legends

Brooks et al., Figure S1

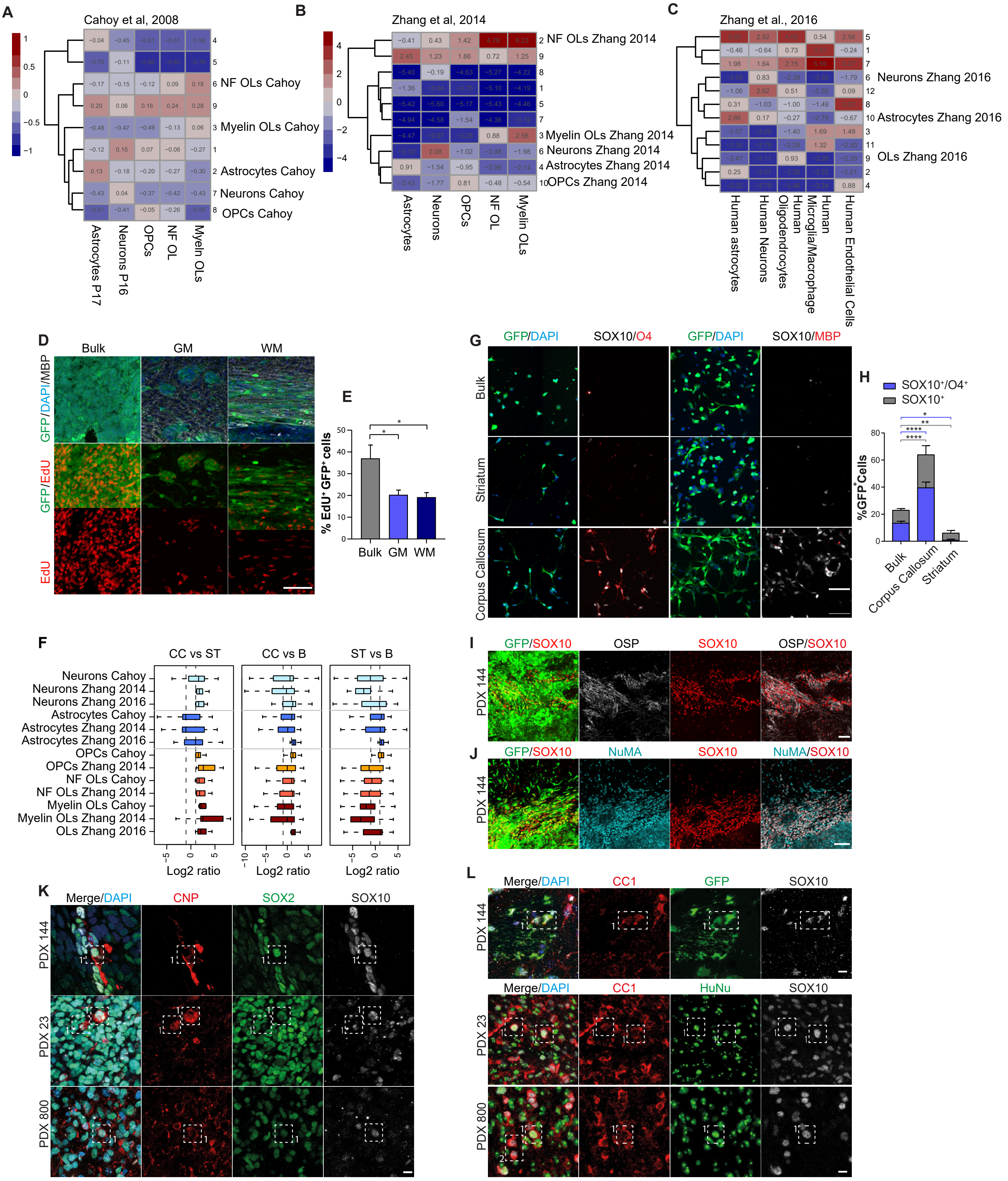

**Figure S1. Derivation of brain cell signatures.**

**A-C**, K-means clustering analysis of median centered expression scores for genes highly variable in published cell-type specific mouse and human brain transcriptomes. Specific clustres were used to define cell type signatures as indicated on the figure (Table S4). **D**, Representative immunofluorescence images and **E**, quantifications of EdU incorporation (red) of $GFP^{+}$ tumour cells of the bulk, grey matter (GM) and white matter (WM) of G144 xenografts. A minimum of 200 cells per xenograft were counted. Mean±SEM, n=3-4 xenografts. *p<0.05. Scale bar=100µm, Two-way ANOVA with Sidak's multiple comparisons test. **F**, Boxplots of DEseq2 xpression ratio for indicated gene signatures in CC vs ST (left), CC vs B (middle) and ST vs B (right) comparisions. The whiskers extend to the most extreme data point, which is no more than 1.5× the interquartile range from the box. Vertical dashed lines mark the 1 and -l x-values (2 fold increase or decrease on a linear scale). **G**, representative images and **H**, quantifications of $GFP^{+}$ tumour cells isolated from indicated tumour regions, acutely cultured and stained for SOX10 (grey) and the pre-oligodendrocyte marker O4 (red) or the myelinating oligodendrocyte marker MBP (red). A minimum of 140 cells per xenograft were counted. Mean±SEM, n=6 xenografts. ****p<0.0001, **p<0.01, *p<0.05. Scale bar=100µm, Two-way ANOVA with Sidak's multiple comparisons test. **I**, high magnification images of SOX10 (grey) and OSP (red) immunofluorescence staining of GFP-labelled G144 tumours. Note disrupted appearance of $OSP^{+}$ myelin fibres. Scale bar=100µm. **J**, high magnification images of NuMA (turquoise), SOX10 (red) and DAPI (blue) immunofluorescence staining of G144 tumours. Scale bar=100µm. **K**, immunofluorescence staining for the immature oligodendrocyte markers CNP (red) and **L**, CC1 (red) in a panel of xenografts. SOX2, GFP or HuNu (green) were used to identify tumour cells as indicated. Dashed square boxes highlight examples of marker positive tumour cells. Scale bar=10µm. Related to Figure 1.

Brooks et al., Figure S2

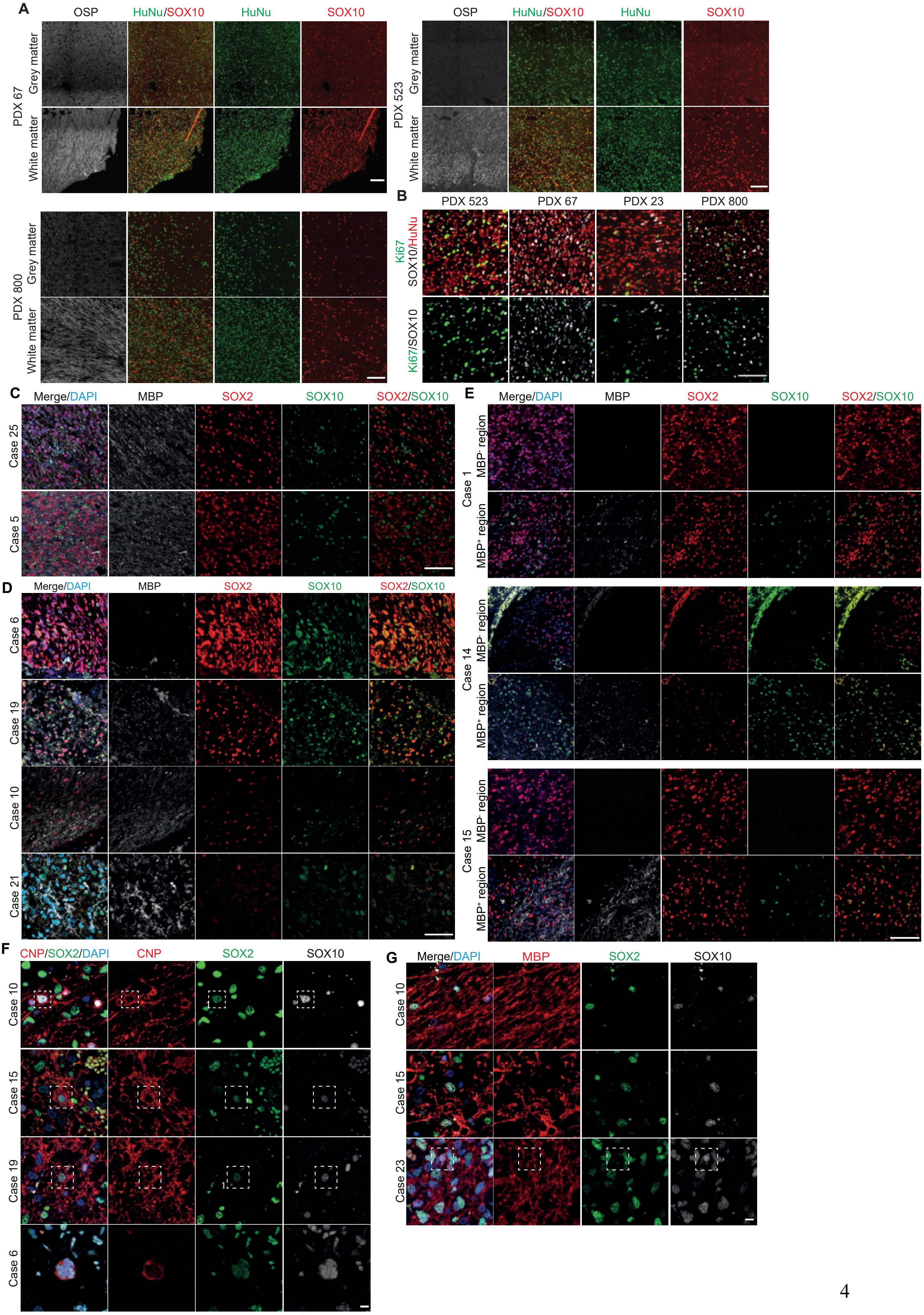

**Figure S2: SOX10 expression in patient-derived xenografts and primary patient tumours. A,** representative immunofluorescence images of $SOX10^{+}$ patient-derived xenografts quantified in Figure 2B (PDX 67, 523, 800) stained for SOX10 (red), OSP (grey) and HuNu (green). Scale bar=100µm. **B**, SOX10 (grey), HuNu (red) and Ki67 (green) immunofluorescence images of PDX tumours quantified in Figure 2C. **C,** MBP (grey), SOX10 (green) and SOX2 (red) immunofluorescence staining of white matter regions of $SOX10^{-}$ patient tumours. No or rare $SOX2^{+}/SOX10^{+}$ endogenous OPCs are observed, confirming that SOX2 can be used to identify tumour cells. Scale bar=100µm. **D,** Representative images of patient tumours stained for SOX2 (red), SOX10 (green), MBP (grey) and DAPI (blue). Scale bar=100µm. **E**, Representative images of patient tumours stained for SOX2 (red), SOX10 (green), MBP (grey) and DAPI (blue) in tumour areas containing ($MBP^{+}$) and devoid of ($MBP^{-}$) white matter. Scale bar=100µm. Note that SOX10 induction is specific to white matter regions. **F**, immunofluorescence staining for the immature oligodendrocyte marker CNP (red) and **G**, the myelinating oligodendrocyte marker MBP (red) in patient material. SOX2 (green) was used to identify tumour cells. Dashed square boxes highlight examples of marker positive tumour cells in F and endogenous oligodendrocytes in G. No $MBP^{+}$ tumour cells are observed amongst $MBP^{+}$ endogenous oligodendrocytes. Scale bar=10µm. Related to Figure 2.

Brooks et al., Figure S3

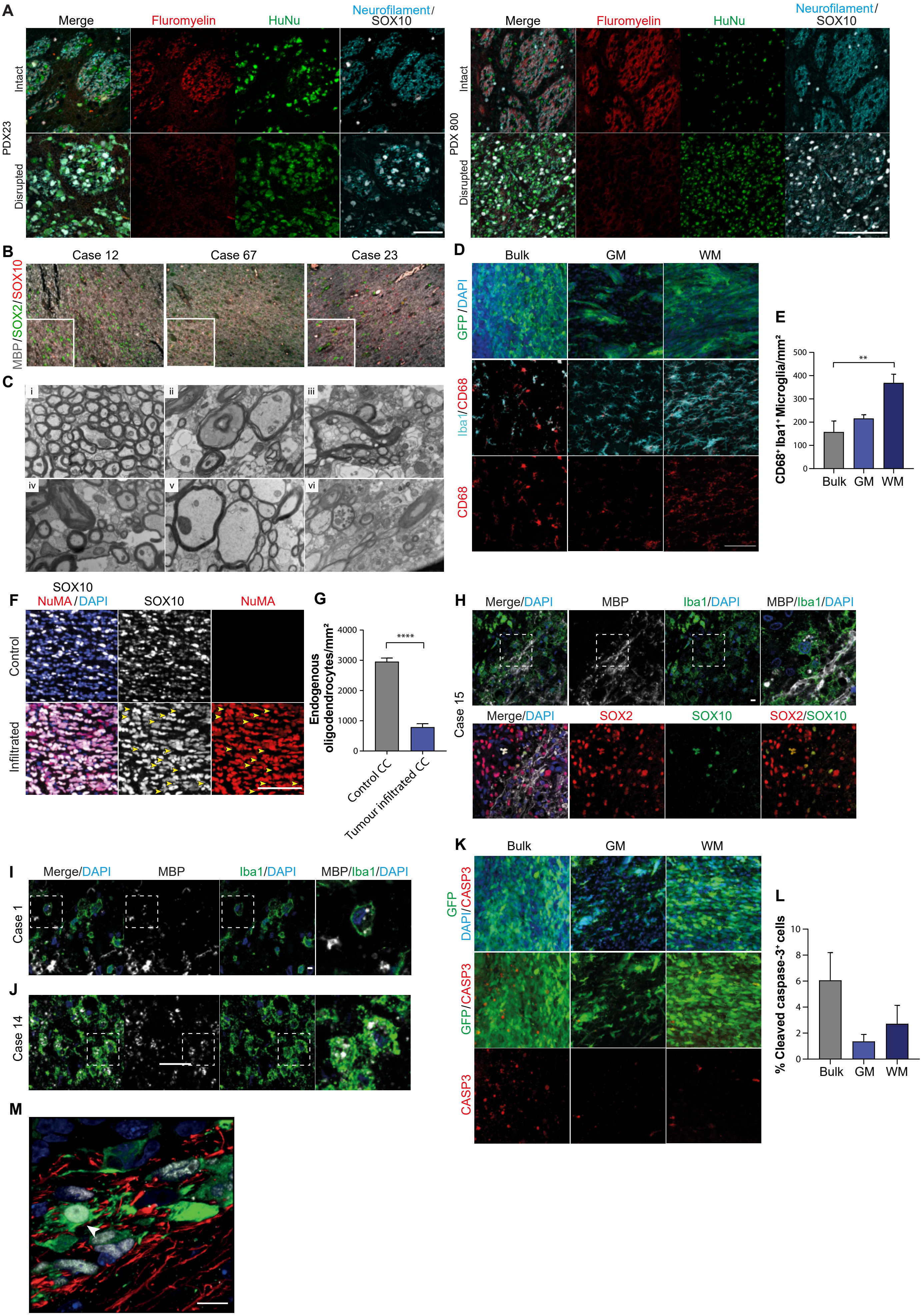

**Figure S3. SOX10 upregulation does not occur in intact myelin.**

**A,** Fluoromyelin (red), SOX10 (grey), Neurofilament (turquoise) and HuNu (green) staining of white matter regions of xenografts quantified in Figure 3B. Scale bar=100µm. **B,** representative images of patient tumours stained for SOX2 (green), SOX10 (red), MBP (grey) and DAPI (blue). Note that the majority of $SOX2^{+}$ tumour cells do not express SOX10 in regions of more intact myelin. Scale bar=50µm. **C**, EM micrographs depicting examples of axonal degeneration and myelin disruption phenotypes observed in tumour-infiltrated white matter: *i* intact contralateral axons, *ii* dark axon, *iii* vacuolised axon, *iv* axon with condensed axoplasm, *v* swollen axon, *vi* axon with enlarged organelles. Scale bar=1µm. **D**, Representative immunofluorescence images and **E**, quantifications of $Iba1^{+}$ (turquoise)/$CD68^{+}$ (red) activated microglia in the bulk, grey matter (GM) and white matter (WM) of $GFP^{+}$ G144 xenografts. A minimum of 140 cells per region per xenograft were counted. Mean±SEM, n=3-4 xenografts. **p<0.01. Scale bar=100µm, Two-way ANOVA with Sidak's multiple comparisons test. **F**, representative immunofluorescence image of the corpus callosum invaded by tumour cells in a terminal G144 xenograft stained for SOX10 in grey, NuMA in red and DAPI in blue. Scale bar=50µm. **G**. quantification of $SOX10^{+}$/$NuMA^{-}$ endogenous oligodendroglia in the images in F. A minimum of 450 cells across 2 ROIs per xenograft were counted. Mean±SEM, n= 2 normal brains, n=5 xenografts **p<0.01. Unpaired two-tailed Student's t test. **H-J**, immunofluorescence images of $Iba1^{+}$ microglia (green) with engulfed MBP myelin debris (grey, dashed box) in tumour infiltrated white matter of patient tumours. For case 15 in H the sequential section was stained for SOX2 (red) and SOX10 (green), confirming that tumour cell differentiation occurs in areas of demyelination (bottom panels). Scale bar=10µm. **K**, representative immunofluorescence images and **L**, quantifications of cleaved $Caspase^{+}$ (red) staining of $GFP^{+}$ (green) tumour cells in the bulk, grey matter (GM) and white matter (WM) regions of GFP-labelled G144 xenografts. A minimum of 200 cells per region per xenograft were counted. Scale bar=100µm Mean±SEM, n=3 xenografts. No significant differences in apoptosis were found. Two-way ANOVA with Sidak's multiple comparisons test. **M**, representative super-resolution image of a demyelinating region within a G144 xenograft. Sections were stained for SOX10 (grey), neurofilament (NF, red). Endogenous GFP fluorescence is in green and nuclei are counterstained with DAPI (blue). Arrowhead indicates an pre-oligodendrocyte-like tumour cell with multiple cellular processes aligned to axons. Scale bar=10µm. Related to Figure 3.

Brooks et al., Figure S4

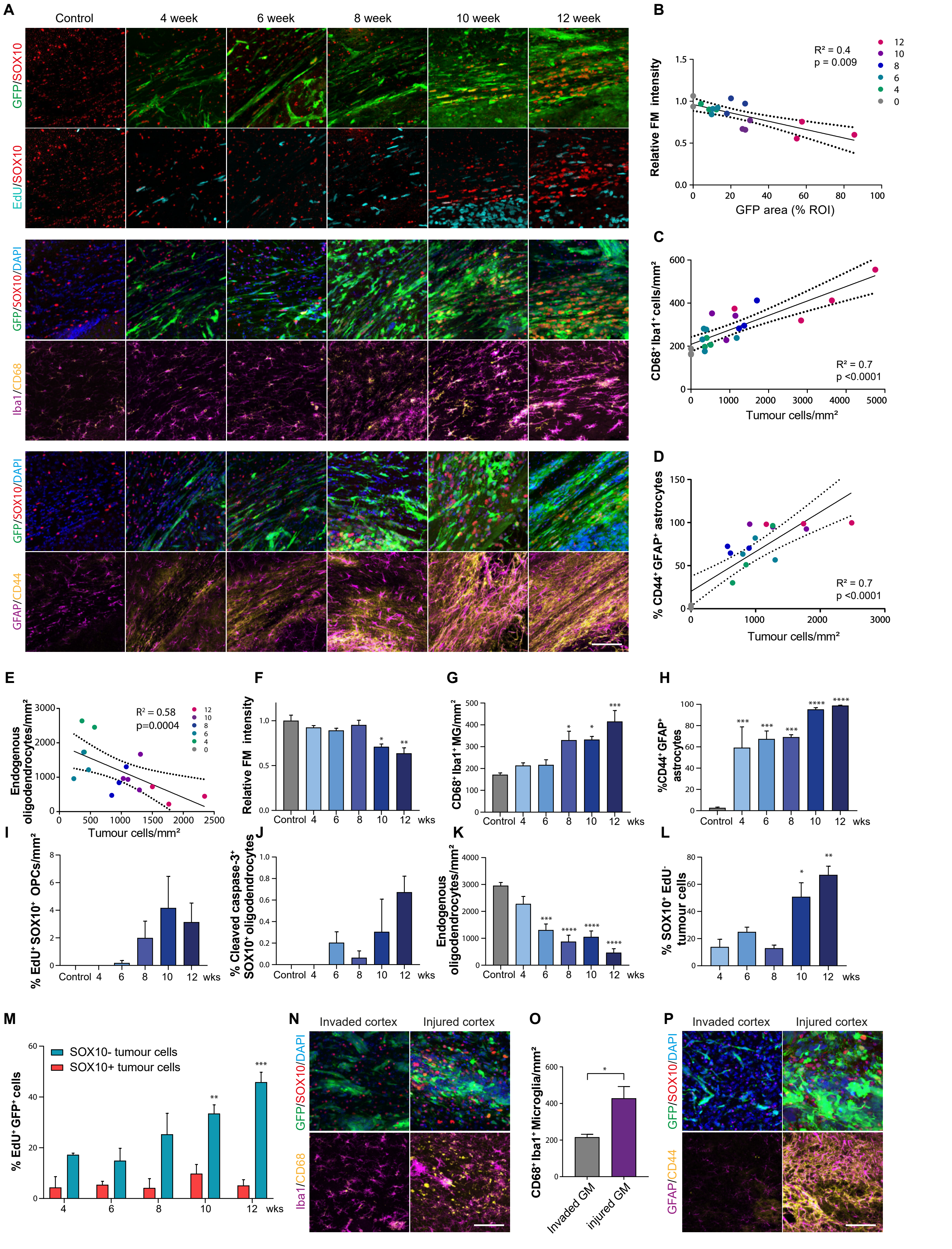

**Figure S4. GBM differentiation is an injury response.**

**A**, time-course analysis of GBM cell infiltration into the corpus callosum in GFP-labelled G144 xenografts collected at indicated time-points (weeks). Representative immunofluorescence images of each time point are shown. All samples were stained for SOX10 (red) and GFP (green) to visualise extent of infiltration and tumour cell differentiation (top panels). In addition, top samples were stained for SOX10 (red) and EdU (turquoise) to assess stability of differentiation, middle samples for Iba1 (magenta) and CD68 (yellow) to assess microglia activation and lower panels for GFAP (magenta) and CD44 (yellow) to examine reactive astrocytes**.** Scale bar=100µm. **B**, quantification of fluoromyelin (FRM) intensity as a function of tumour density. Dots indicate individual xenografts colour-coded by colletion time-point. n=3 tumours/time point, n=2 intact control brains. The coefficient of determination is indicated. **C**, quantification of the number activated microglia, **D**, reactive astrocytes and **E** endogenous oligodendrocytes in the corpus callosum as a function of number of invaded tumour cells. Dots indicate mean cell density in individual xenografts colour-coded by time. A minimum ROI area of 300µm was analysed. The coefficient of determination and p value are indicated. **F-L**, time-course analysis of tumour cell differentiation and glial response during tumour infiltration into the corpus callosum in G144 xenografts. Shown is the quantification of indicates cell types over time. A minimum ROI area of 300µm was analysed within intact or disrupted white matter. Mean±SEM, n=3 xenografts. ****p<0.0001, ***p<0.001, **p<0.01, *p<0.05, ANOVA with Dunnett's multiple comparison tests. **M**, quantification of the percentage of proliferative ($EdU^+$) differentiated $SOX10^+$ and non-differentiated $SOX10^-$ $GFP^+$ G144 cells in the time-course experiments shown in A. Note that the percentage of $SOX10^+/EdU^+$ remained constant indicative of stable differentiation. A minimum ROI area of 300µm was analysed. Mean±SEM, n=3 xenografts per time point. ***p<0.001, **p<0.01. Two-way ANOVA with Sidak's multiple comparisons test. **N**, representative immunofluorescence images and **O**, quantifications of SOX10 (red), Iba1 (magenta), CD68 (yellow) and DAPI (blue) immunofluorescence staining of $GFP^+$ G144 cells directly injected (injured CTX) or invaded into the dense myelin of inner cortex from the tumour bulk (invaded CTX). Scale bar=100μm. A minimum of 90 cells per xenograft were counted. Mean±SEM, n=3 xenografts per group. *p<0.05. Unpaired two-tailed Student's t test. **P**, SOX10 (red), GFAP (magenta), CD44 (yellow) and DAPI (blue) immunofluorescence staining of $GFP^+$ G144 cells directly injected (injured CTX) or invaded into the dense myelin of inner cortex from the tumour bulk (invaded CTX). Note that injection into the inner cortex causes glial activation consistent with an injury response. Scale bar=100μm. Related to Figure 3.

Brooks et al., Figure S5

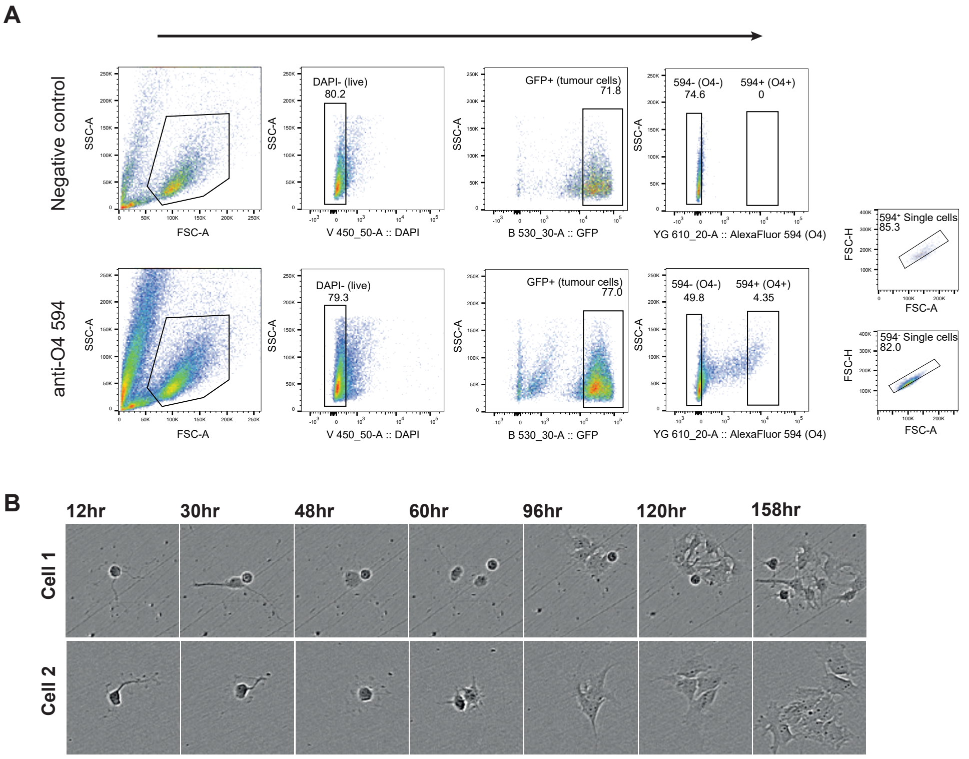

**Figure S5. Differentiation is reversed upon removal from the white matter. A**, representative FACS plots of the purification of O4$^+$/GFP$^+$ G144 cells from CC and B regions of xenografts. Unstained GFP$^+$ G144 cells from the same regions were used for gating. **B**, representative phase contrast still images taken from videos of O4$^+$ G144 cells acutely FACS-sorted from the B and CC regions of primary xenografts and seeded in neural stem cell conditions for 7 days. Related to Figure 4.

Brooks et al., Figure S6

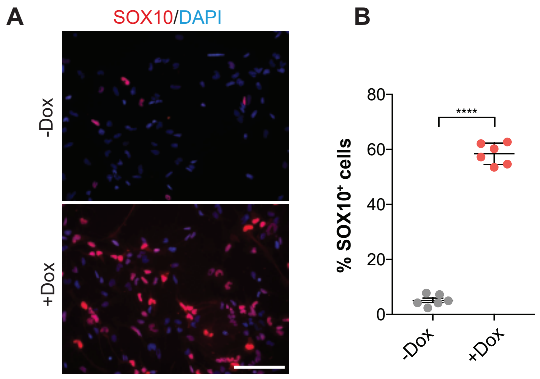

**Figure S6. Efficacy of SOX10 overexpression.**

**A**, representative images of SOX10 (red) and DAPI (blue) fluorescence staining of G144 cultures before (Dox-) and after (Dox+) SOX10 induction with doxycycline for 48h. Scale bar=100µm. **B**, quantification of percentage of SOX10 expressing cells in cultures from a. A minimum of 90 cells were counted per group. Mean±SEM, n=2 independent cultures each on triplicate coverslips. Related to Figure 5.

Brooks et al., Figure S7

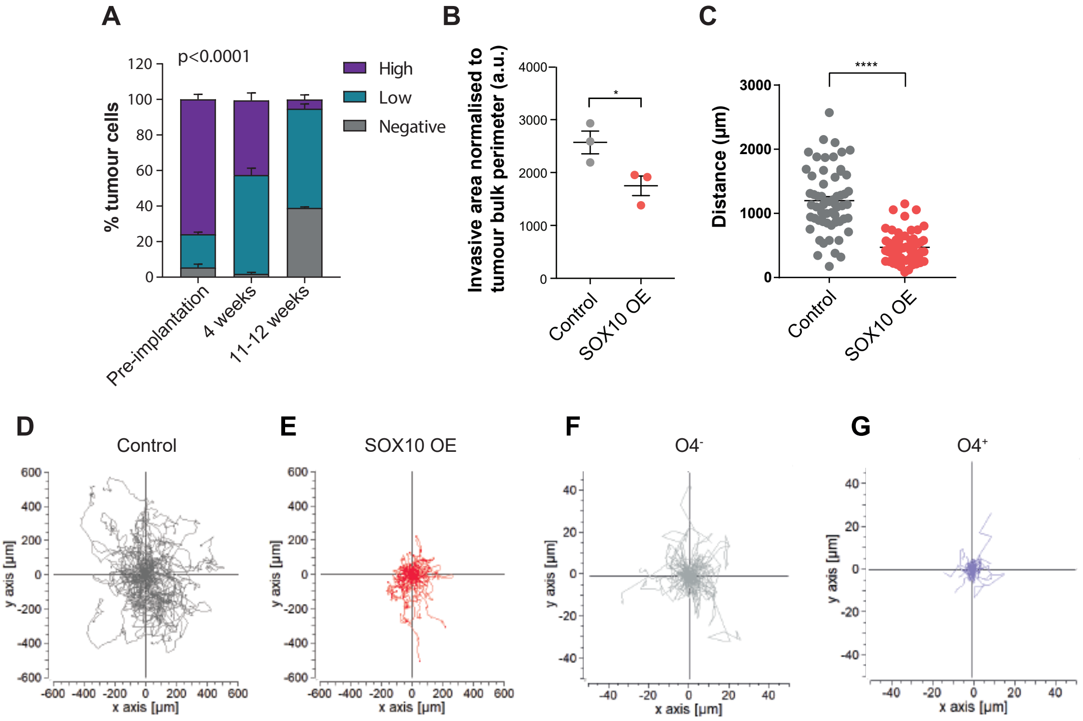

**Figure S7. SOX10 upregulation reduces tumour cell motility.**

**A**, quantification of percentages of $SOX10^-$, SOX10low and SOX10high tumour cells in G144 SOX10 OE xenografts. Quantifications were carried out *in vitro* before tumour implantation (pre-implantation) or *in vivo* at 4 or 11-12 weeks post-implantation in grey matter regions of xenografts, as indicated. A minimum of 250 cells per group were counted. Mean±SEM, n=3 cultures or xenografts per group. **** p<0.0001. Two-way ANOVA. **B**, quantification of area occupied by invasive tumour cells in Control and SOX10 OE tumours shown in Figure 6E. Quantifications were performed across 2 sections per mouse. Mean±SEM, n=3 mice/group. Unpaired two-tailed Student's t test. **C**, quantification of migrated distance of individual G144 cells in Control and SOX10 OE G144 cultures. Each dot represents a cell. Mean±SEM, n=60 cells per group pooled from 3 independent transductions. ****p<0.0001. Unpaired Student's t-tes. **D**, **E** traces of individual migrating cells from C aligned to the same point of origin. **F**, **G**, Traces of individual $O4^+$ and $O4^-$ tumour cells from Figure 6G, which were acutely isolated from the corpus callosum region of G144 xenografts and cultured for 72h in neural stem cell conditions. Traces were realigned to the same point of origin. Related to Figure 6..

Brooks et al., Figure S8

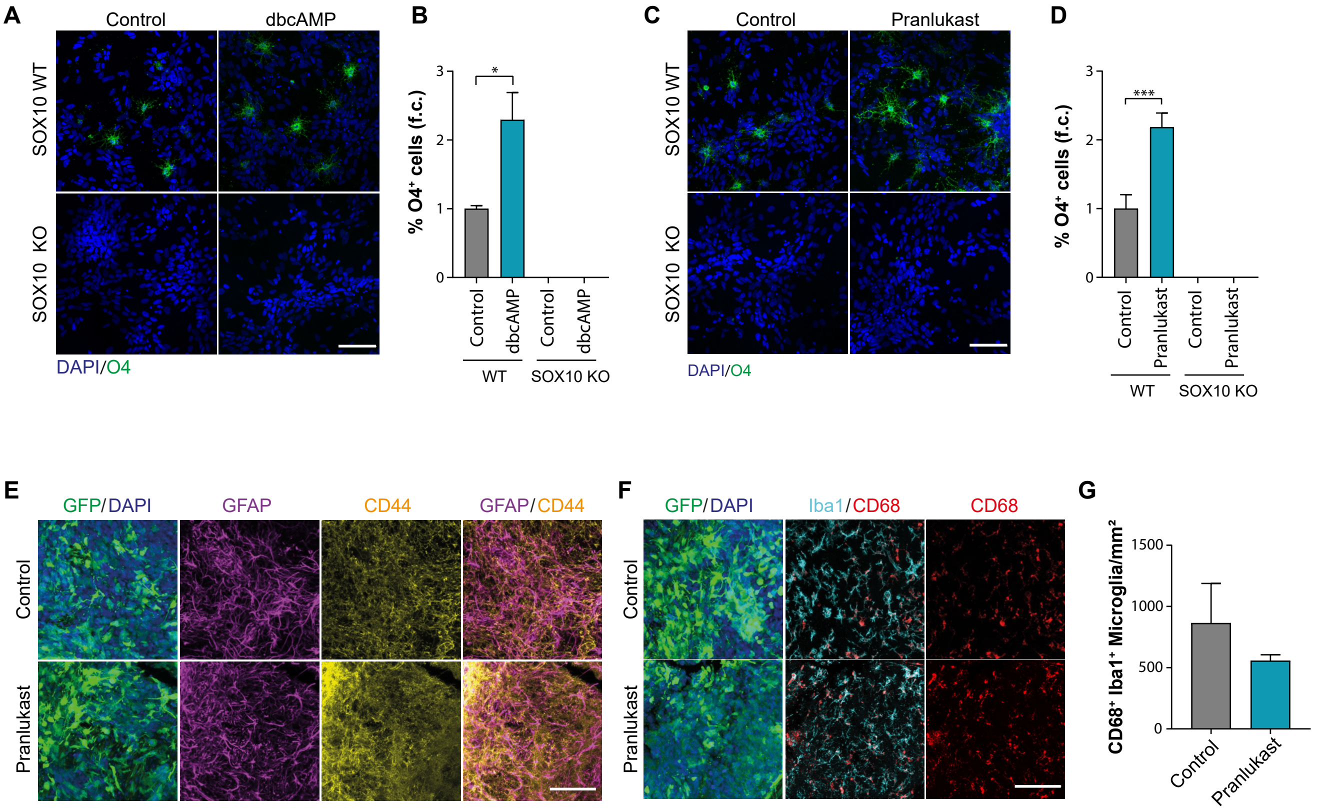

**Figure S8. Myelination-promoting drugs induce GBM differentiation via SOX10.**

**A**, representative immunofluorescence images and **B** quantifications of control and SOX10 knock-out G144 cultures untreated or treated with db-cAMP for 14 days in the absence of growth factors and stained for $O4^+$ (green). **C**, representative immunofluorescence images and **D** quantifications of control and SOX10 knock-out G144 cultures untreated or treated with Pranlukast for 14 days in the absence of growth factors and stained for $O4^+$ (green). A minimum of 1000 cells cells across duplicate coverslips was counted per biological repeat. Mean±SEM, n=5 independent cultures per group *$p<0.05$. Unpaired two-tailed Student's t test. **E**, GFAP (turquoise), CD44 (red) and DAPI (blue) immunofluorescence staining of activated astrocytes in $GFP^+$ G144 tumours treated with saline or Pranlukast. **F**, representative immunofluorescence images and **G**, quantifications of $Iba1^+$ (magenta)/$CD68^+$ (yellow) activated microglia staining in $GFP^+$ G144 tumours treated with saline or Pranlukast Scale bar=100µm. A minimum of 400 cells per xenograft were counted. Mean±SEM, n=3 xenografts per group. *$p<0.05$. Unpaired two-tailed Student's t test. Note that Pranlukast did not affect endogenous glial. Scale bar=100µm. Related to Figure 8.

**Video S1**

3D animation of a representative immunofluorescence image of a GFP-labelled G144 tumour stained for CD68 (red) and MBP (grey) showing the presence of microglia with engulfed myelin debris in infiltrated white matter.

**Video S2 and 3**

Time-lapse imaging of $O4^+$ tumour cells FAC-sorted from the corpus callosum of G144 tumours and cultured in stem cell media for 7 days. Two examples of cells losing oligodendrocyte morphology and re-entering the cell-cycle are shown.
